# Effect of vaccinating health care workers to control Ebola virus disease: a modelling analysis of outbreak data

**DOI:** 10.1101/113506

**Authors:** Alexis Robert, Anton Camacho, W. John Edmunds, Marc Baguelin, Jean-Jacques Muyembe Tamfum, Alicia Rosello, Sakoba Kéïta, Rosalind M. Eggo

## Abstract

**Background:** Health care workers (HCW) are at risk of infection during Ebola virus disease outbreaks and therefore may be targeted for vaccination before or during outbreaks. The effect of these strategies depends on the role of HCW in transmission which is understudied.

**Methods:** To evaluate the effect of HCW-targeted or community vaccination strategies, we used a transmission model to explore the relative contribution of HCW and the community to transmission. We calibrated the model to data from multiple Ebola outbreaks. We quantified the impact of ahead-of-time HCW-targeted strategies, and reactive HCW and community vaccination.

**Results:** We found that for some outbreaks (we call “type 1”) HCW amplified transmission both to other HCW and the community, and in these outbreaks prophylactic vaccination of HCW decreased outbreak size. Reactive vaccination strategies had little effect because type 1 outbreaks ended quickly. However, in outbreaks with longer time courses (“type 2 outbreaks”), reactive community vaccination decreased the number of cases, with or without prophylactic HCW-targeted vaccination. For both outbreak types, we found that ahead-of-time HCW-targeted strategies had an impact at coverage of 30%.

**Conclusions:** The optimal vaccine strategy depends on the dynamics of the outbreak and the impact of other interventions on transmission. Although we will not know the characteristics of a new outbreak, ahead-of-time HCW-targeted vaccination can decrease the total outbreak size, even at low vaccine coverage.

**summary:** Targeting health care workers for Ebola virus disease vaccination can decrease the size of outbreaks, and the number of health care workers infected. The impact of these strategies decrease depends on timing, coverage, and the dynamics of the outbreak.

## Background

Sub-Saharan Africa has experienced more than 26 Ebola virus disease (EVD) outbreaks since 1976. The largest of these, the 2013-16 West African outbreak, resulted in more than 28,000 cases in Liberia, Sierra Leone and Guinea [1]. In this and in many other EVD outbreaks there was higher incidence among health care workers (HCW) than in the general population [2–6].

HCW are at high risk of infection from patients (especially before introduction of personal protective equipment (PPE) [7]) due to their frequent and close contact with them, for example during surgical procedures. Infected HCW may in turn transmit the infection to coworkers or other patients [7]. Nosocomial EVD outbreaks can also spread to the wider community.

Mathematical modelling can provide insight into key epidemiological drivers and anticipate the effect of control measures [8–14]. Few models have investigated the role of HCW in transmission, despite strong evidence of their importance during outbreaks [2–4,15]. Understanding the role of HCW in transmission is crucial to appropriately assess the potential benefit of HCW-targeted vaccination, so that appropriate policy decisions can be made once an EVD vaccine is licensed.

To evaluate HCW-targeted vaccination strategies, we used a mechanistic transmission model to separately estimate the effect of HCW and community members. By calibrating this model to observed patterns of infections from prior EVD outbreaks, we could compare the effect of different vaccination strategies.

## Methods

### Outbreak Data

Information on the occupation of cases is rarely available [16], although this information is critical to determining the role of HCW in transmission. We used data for outbreaks where we could find occupation (HCW or not) of cases. This resulted in twelve timeseries drawn from local outbreaks during the West African epidemic, and from the large 1995 Kikwit outbreak in the Democratic Republic of Congo (DRC) (Supplementary Section 1). All twelve timeseries are provided in the supplement, and five (for brevity) are shown in Figure 1. We noted that the number, timing, and dynamics of HCW infections during these outbreaks were not consistent, and we therefore used the dynamics of HCW and community infections to classify the outbreaks into types.

**Figure 1.**
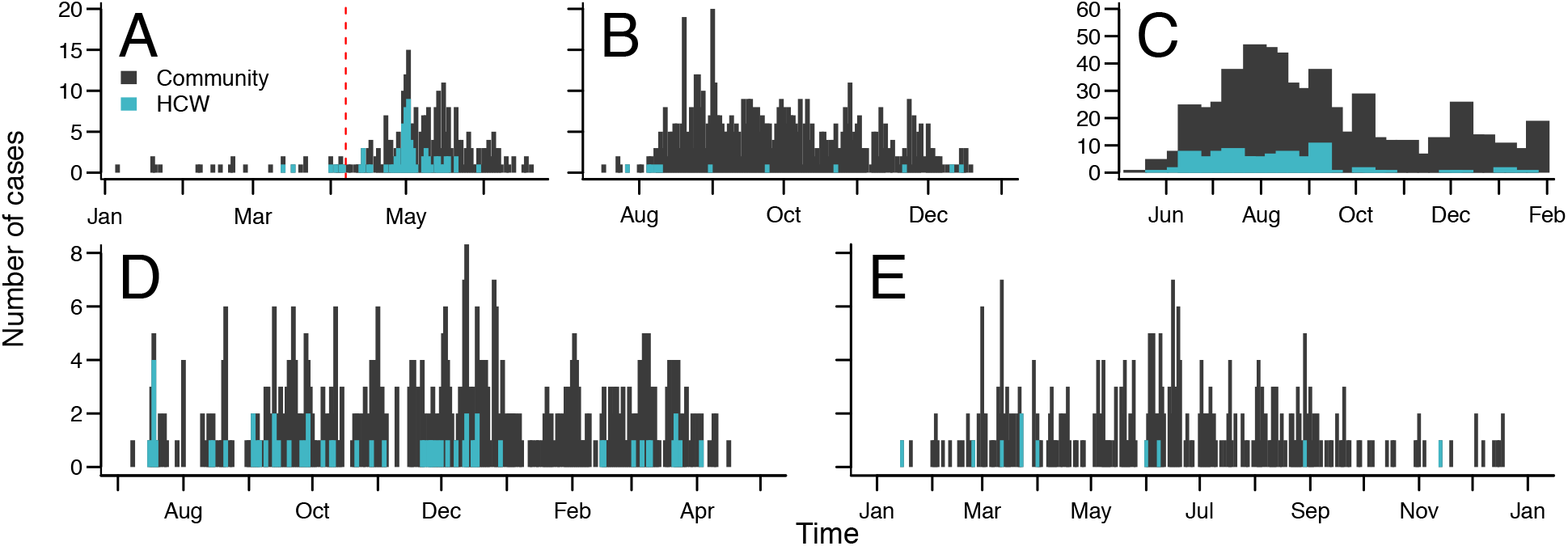
Incidence time series of EVD onset in five of the outbreaks used. Daily incidence in A) Kikwit (DRC 1995), B) Macenta (Guinea 2014), D) Conakry (Guinea, 2014-2015) and E) Gueckedou (Guinea, 2014); C) Weekly incidence in Kenema (Sierra Leone, 2014). Bong County, Liberia is not shown here because the data are infrequently reported making incidence estimation difficult. Cumulative statistics are shown in Figure 2 for Bong County.

### Classification of EVD outbreaks into two types

We compared the HCW infection dynamics of the outbreaks and determined that they fell loosely into two types. The distinguishing characteristics we used were grouped into four categories: i) the proportion of HCW and community infected through time; ii) the shape and timing of the cumulative distribution of HCW infections; iii) the weekly proportion of HCW infected; and iv) the total size of the outbreak in both number of cases and duration (Supplementary section S5). Data from Kikwit (1995, DRC) were uniquely detailed and therefore provided an opportunity to quantify the role of HCW in transmission by fitting a mechanistic model. According to our classification, Kikwit was a “type 1” outbreak (described in Results), and therefore we also calibrated the transmission model to the other observed outbreak pattern: “type 2”.

### Kikwit outbreak data

These data contained epidemiologically-inferred links between cases, the occupations of both infectors and infectees, and daily granularity of symptom onset (Supplementary Section 2). The outbreak started with infrequent cases in rural areas before introduction to Kikwit General Hospital on April 7^th^ (Figure 1a) [6]. On May 2^nd^ a haemorrhagic fever was diagnosed, on May 8^th^ this was confirmed as EVD, and on May 10^th^ international assistance was initialised. Further control measures started on May 12^th^ (Supplementary Section S2). The final case died on July 16^th^, resulting in 317 cases reported, 248 deaths, and a case-fatality ratio of 78% [17].

For each case we used their occupation; the occupation of their likely infector (obtained by real-time epidemiological investigation); date of onset; and date of recovery or death. There were some missing data in each field (Supplementary Section S2). We censored cases with date of onset before April 7^th^, when the first case was admitted to Kikwit General Hospital, which gave 284 cases, of whom 73 were HCW (26%). A likely infector was available for 191 cases.

### Transmission Model

We developed a deterministic compartmental model of EVD transmission stratified by occupation, where individuals were either HCW (*h*) or community (*c*) (Figure 3). On infection, cases left the susceptible compartment (*S*), and entered latent infection (*E)*. The duration in *E* was drawn from an Erlang distribution with shape 2 and mean *ϵ^-1^* [18].

Following *E*, individuals became infectious and symptomatic (*I*), and recovered (*R*) or died (*D*). We did not have information about funeral transmission in Kikwit, and thus we considered that all transmission events occurred from the *I* compartment (Supplementary Section S3). For each occupation we defined four time-dependent transmission rates: *β_t,ij_*, where infectious, *i*, and susceptible, *j*, are either *c* or *h*. We assumed a single introduction to each population, and no population movement during the outbreak.

To account for the effect of changing transmission rates, resulting from control measures such as the arrival of PPE for HCWs, opening of isolation wards, and population awareness of EVD, we used flexible time-dependent sigmoid functions for the transmissibility parameters, *β_tij_*, for which we estimated the parameters (Supplementary Section S3). The force of infection was *λ_t,ij_* = *β_t,ij_*(*l_i_/N_j_*), where *N_j_* was the population size of HCW (900) or community members (200,000) in Kikwit in 1995 (Supplementary Section S2).

### Model fitting for Kikwit outbreak

We fitted the model to six time series of onset dates stratified by case and infector occupation, and two time series of deaths stratified by case occupation. We used a negative binomial likelihood and Bayesian methods [19, 20] and non-informative priors (Supplementary section S4). We calculated the estimated reproduction number by occupation of infector and infectee (*R_ij_*), and the net reproduction number (*R_n_*) using the next generation matrix (Supplementary section S3).

### Calibration of type 2 outbreak scenario

Data from available type 2 outbreaks did not include links between cases, and therefore were too incomplete to fit the same model framework. Observed type 2 outbreaks were characterised by a longer time period of HCW infections with no early rapid increase, and the epidemics were longer and larger in both occupation groups (see Results and Figure 2). We used published evidence to calibrate the type 2 scenario. We used initial HCW and community reproduction numbers from a large study of transmission in Guinea [21], which found low HCW-related transmission, and higher community-related transmission.

**Figure 2.**
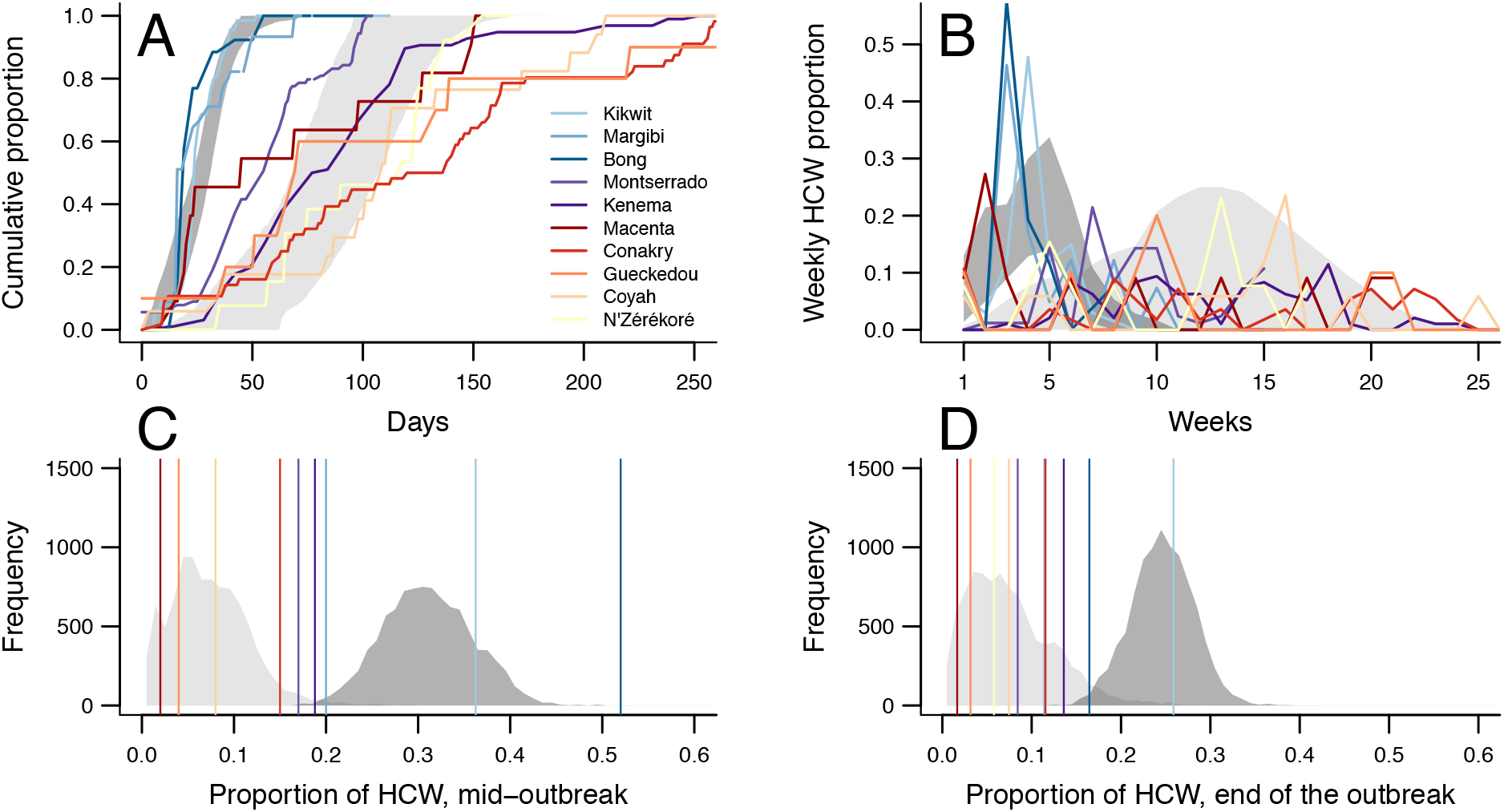
Classification of EVD outbreaks into two broad types. Dark grey marks the 95% CI in type 1 outbreak simulations, light grey areas correspond to 95% CI for type 2 outbreak simulations. Each colour corresponds to an outbreak, we used blue shades for type 1 outbreaks and red/purple shades for type 2. A) Cumulative proportion of HCW among the total number of HCW infected through time. B) Weekly proportion of HCW among the total number of HCW infected through time. C) Proportion of HCW among all cases infected at mid-outbreak. D) Proportion of HCW among all cases infected at the end of the outbreak Vertical lines in Figure C and D correspond to the value for each outbreak.

Contemporaneous analyses of the West African outbreak suggested a pattern of initially sustained transmission in the community, followed by a slow decline in transmission [22–25]. Therefore, we calibrated the parameters of the sigmoid functions to give a slow decrease in *β_t,ij_*. We computed the four reproduction numbers between each occupation group using these published estimates for the overall reproduction numbers. To fully quantify the uncertainty, we used the parameter uncertainty from fitting the Kikwit data. The modifications we made to the transmission parameters (Table 2) resulted in simulated outbreaks with higher community reproduction number and slower transmission decrease in type 2 compared with type 1 outbreaks. We kept the same parameters for population size, number of HCW, and reporting fractions as in Kikwit, which allowed direct comparison of type 1 and 2 scenarios, and therefore the impact of vaccination.

**Table 1.**
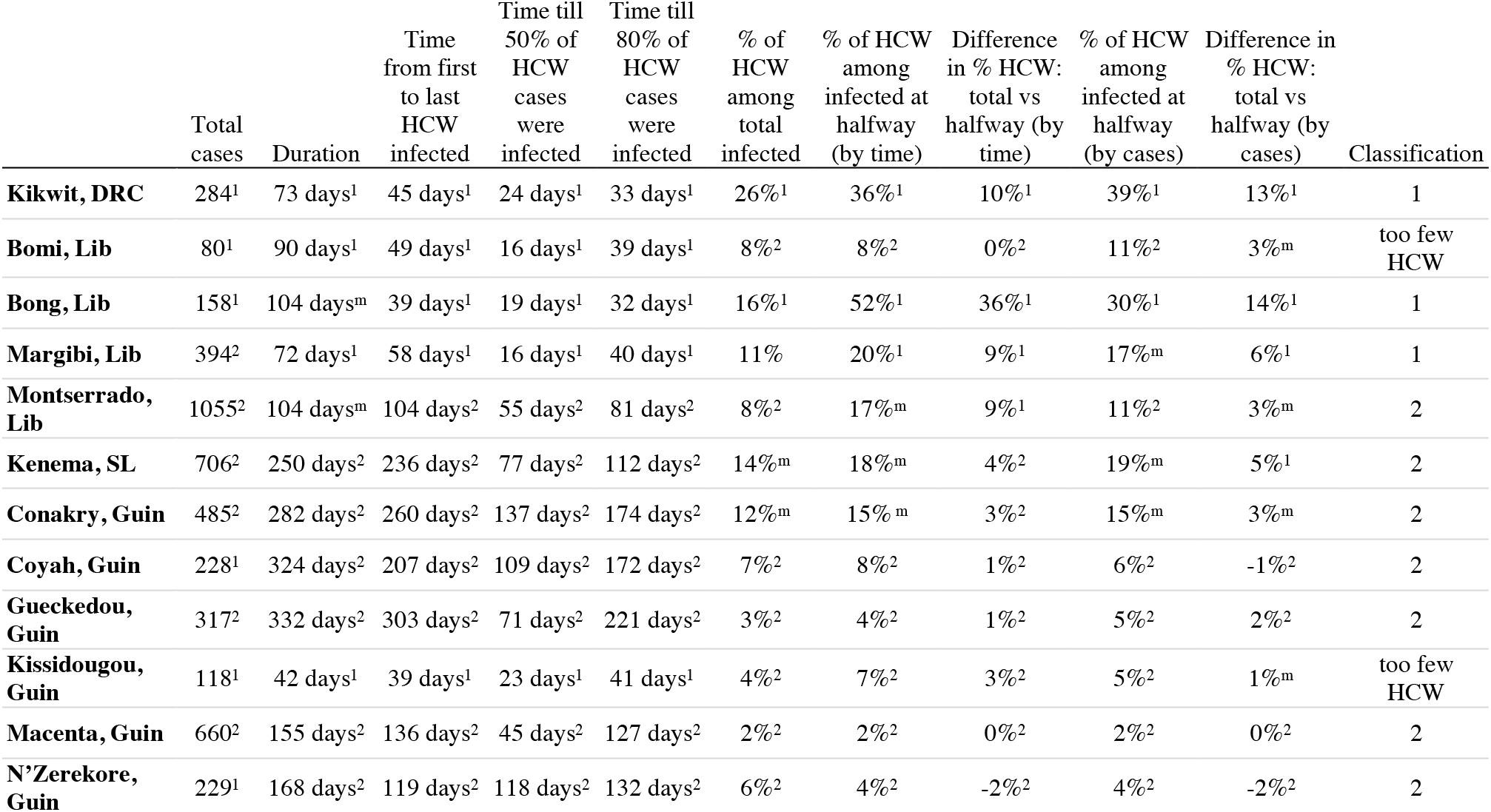
Characteristics of the outbreaks. ^1^marks type 1, ^2^marks type 2, and ^m^ a mixed or intermediate value. Thresholds are as follows: Total cases (1=1-250, mixed=250-300, 2=300+), Duration (1-99, 100-150, 150+), Time till 50% (1-24, 25-49, 50+), Time till 80% (1-44, 45-70, 70+), % HCW among total (15+, 10-14, 1-10), % at halfway by time (20+, 15-19, 0-14), Difference in % at halfway by time (7.5+, 5-7.49, 0-4), % at halfway by cases (20+, 10-19, 1-10), Difference in % at halfway by cases (5+, 3-4, <2). These values, together with the graphs showing the dynamics of infection were used to classify the outbreaks into two broad types.

|  | Total cases | Duration | Time from first to last HCW infected | Time till 50% of HCW cases were infected | Time till 80% of HCW cases were infected | % of HCW among total infected | % of HCW among infected at halfway (by time) | Difference in % HCW: total vs halfway (by time) | % of HCW among infected at halfway (by cases) | Difference in % HCW: total vs halfway (by cases) | Classification |
| --- | --- | --- | --- | --- | --- | --- | --- | --- | --- | --- | --- |
| **Kikwit, DRC** | 284<sup>1</sup> | 73 days<sup>1</sup> | 45 days<sup>1</sup> | 24 days<sup>1</sup> | 33 days<sup>1</sup> | 26%<sup>1</sup> | 36%<sup>1</sup> | 10%<sup>1</sup> | 39%<sup>1</sup> | 13%<sup>1</sup> | 1 |
| **Bomi, Lib** | 80<sup>1</sup> | 90 days<sup>1</sup> | 49 days<sup>1</sup> | 16 days<sup>1</sup> | 39 days<sup>1</sup> | 8%<sup>2</sup> | 8%<sup>2</sup> | 0%<sup>2</sup> | 11%<sup>2</sup> | 3%<sup>m</sup> | too few HCW |
| **Bong, Lib** | 158<sup>1</sup> | 104 days<sup>m</sup> | 39 days<sup>1</sup> | 19 days<sup>1</sup> | 32 days<sup>1</sup> | 16%<sup>1</sup> | 52%<sup>1</sup> | 36%<sup>1</sup> | 30%<sup>1</sup> | 14%<sup>1</sup> | 1 |
| **Margibi, Lib** | 394<sup>2</sup> | 72 days<sup>1</sup> | 58 days<sup>1</sup> | 16 days<sup>1</sup> | 40 days<sup>1</sup> | 11% | 20%<sup>1</sup> | 9%<sup>1</sup> | 17%<sup>m</sup> | 6%<sup>1</sup> | 1 |
| **Montserrado, Lib** | 1055<sup>2</sup> | 104 days<sup>m</sup> | 104 days<sup>2</sup> | 55 days<sup>2</sup> | 81 days<sup>2</sup> | 8%<sup>2</sup> | 17%<sup>m</sup> | 9%<sup>1</sup> | 11%<sup>2</sup> | 3%<sup>m</sup> | 2 |
| **Kenema, SL** | 706<sup>2</sup> | 250 days<sup>2</sup> | 236 days<sup>2</sup> | 77 days<sup>2</sup> | 112 days<sup>2</sup> | 14%<sup>m</sup> | 18%<sup>m</sup> | 4%<sup>2</sup> | 19%<sup>m</sup> | 5%<sup>1</sup> | 2 |
| **Conakry, Guin** | 485<sup>2</sup> | 282 days<sup>2</sup> | 260 days<sup>2</sup> | 137 days<sup>2</sup> | 174 days<sup>2</sup> | 12%<sup>m</sup> | 15%<sup>m</sup> | 3%<sup>2</sup> | 15%<sup>m</sup> | 3%<sup>m</sup> | 2 |
| **Coyah, Guin** | 228<sup>1</sup> | 324 days<sup>2</sup> | 207 days<sup>2</sup> | 109 days<sup>2</sup> | 172 days<sup>2</sup> | 7%<sup>2</sup> | 8%<sup>2</sup> | 1%<sup>2</sup> | 6%<sup>2</sup> | -1%<sup>2</sup> | 2 |
| **Gueckedou, Guin** | 317<sup>2</sup> | 332 days<sup>2</sup> | 303 days<sup>2</sup> | 71 days<sup>2</sup> | 221 days<sup>2</sup> | 3%<sup>2</sup> | 4%<sup>2</sup> | 1%<sup>2</sup> | 5%<sup>2</sup> | 2%<sup>2</sup> | 2 |
| **Kissidougou, Guin** | 118<sup>1</sup> | 42 days<sup>1</sup> | 39 days<sup>1</sup> | 23 days<sup>1</sup> | 41 days<sup>1</sup> | 4%<sup>2</sup> | 7%<sup>2</sup> | 3%<sup>2</sup> | 5%<sup>2</sup> | 1%<sup>m</sup> | too few HCW |
| **Macenta, Guin** | 660<sup>2</sup> | 155 days<sup>2</sup> | 136 days<sup>2</sup> | 45 days<sup>2</sup> | 127 days<sup>2</sup> | 2%<sup>2</sup> | 2%<sup>2</sup> | 0%<sup>2</sup> | 2%<sup>2</sup> | 0%<sup>2</sup> | 2 |
| **N'Zerekore, Guin** | 229<sup>1</sup> | 168 days<sup>2</sup> | 119 days<sup>2</sup> | 118 days<sup>2</sup> | 132 days<sup>2</sup> | 6%<sup>2</sup> | 4%<sup>2</sup> | -2%<sup>2</sup> | 4%<sup>2</sup> | -2%<sup>2</sup> | 2 |

**Table 2.** Values of the reproduction number, time of change in transmission, and shape of the decrease in transmission. Type 1 values are inferred from fitting the model to the Kikwit data. Type 2 values are used as described in Methods based on values in [21] to simulate outbreaks. Mean values and 95% CIs are given. Comparison of the *R*_0_ trajectories is given in Supplementary Figure 5.2.

|  |  | HCW to<br>HCW | Community<br>to HCW | HCW to<br>community | Community<br>to community | Overall |
| --- | --- | --- | --- | --- | --- | --- |
| Type 1<br>scenario | $R_0$ | 2.68<br>(1.83-4.01) | 0.16<br>(0.05-0.37) | 3.99<br>(2.47-6.29) | 0.70<br>(0.53-0.95) | 2.98<br>(2.11-4.36) |
| | $T_{change}$<br>(days) | $T_h = 30$ (27-35) | $T_h$ | $T_h$ | $T_c = 55$ (50-62) | |
| | shape | $\alpha_h = 2.20$<br>(0.23-4.83) | $\alpha_h$ | $\alpha_h$ | $\alpha_c = 2.49$<br>(0.21-4.86) | |
| Type 2<br>scenario | $R_0$ | 0.16<br>(0.05-0.37) | 0.16<br>(0.05-0.37) | 0.53<br>(0.24-0.90) | 2.11<br>(1.94-2.34) | 2.15<br>(1.99-2.37) |
| | $T_{change}$<br>(days) | $T_c = 125$<br>(118-133) | $T_c$ | $T_c$ | $T_c$ | |
| | shape | $\alpha_c = 0.05$ | $\alpha_c$ | $\alpha_c$ | $\alpha_c$ | |

### Simulation of vaccination

We extended the model to include vaccination of HCW and community and compared the impact of eight vaccination strategies (Table 3). We sampled 600 parameter sets from the joint posterior distribution and generated 15 stochastic simulations for each. We compared the number of cases and the time to extinction (0 individuals in *E* or *I*) to the baseline scenario without vaccination for each parameter set and random number seed. We used the parameter uncertainty inferred from the Kikwit data for both type 1 and type 2 outbreaks, and report 95% credible intervals in the text.

**Table 3.** Vaccination strategies tested in this analysis. We considered the following: i) reactive mass vaccination of the population, prioritising HCW, with a prime-boost vaccine (strategy a), or a single dose vaccine (strategy b); ii) ahead-of-time vaccination of HCW, with three levels of coverage: 10% (strategy c), 30% (strategy d), or 50% (strategy e); iii) combined strategies of ahead-of-time vaccination of HCW at three levels of coverage plus reactive mass vaccination of remaining HCW and the community (strategies f, g, and h). We selected values of coverage that were realistic given high HCW turnover in recently affected countries [26] [27], and the possibility of waning of protection.

| Strategy | HCW ahead-of-time coverage | HCW reactive coverage | Community reactive coverage | Vaccine |
| --- | --- | --- | --- | --- |
| **a** | 0% | 100% | 70% | Prime-boost |
| **b** | 0% | 100% | 70% | Single dose |
| **c** | 10% |  |  | - |
| **d** | 30% |  |  | - |
| **e** | 50% |  |  | - |
| **f** | 10% | 100% | 70% | Single dose |
| **g** | 30% | 100% | 70% | Single dose |
| **h** | 50% | 100% | 70% | Single dose |

We simulated ahead-of-time HCW vaccine coverage values of 50%, 30%, and 10%. These values reflect the high turnover of HCW in recently affected countries [26, 27], and the possibility that protection could wane. Vaccine coverage should be interpreted as effective levels of coverage: 30% coverage is equivalent to 100% vaccination of HCW and waning to 30% protection, or as 30% vaccination and 100% protection.

Vaccination reduced susceptibility to infection (Figure 3). For single-dose vaccine, efficacy was 90%, and protection was reached after one week. For prime-boost vaccine, efficacy was 90, where 80% was reached one week after prime, and boost was 28 days after prime. We simulated vaccination of 1500 people per day, which was an operational maximum suggested by field teams. Reactive vaccination started with unvaccinated HCW and continued until all HCW and 70% of community members were vaccinated, at the same rate for single dose and prime-boost vaccination.

**Figure 3.**
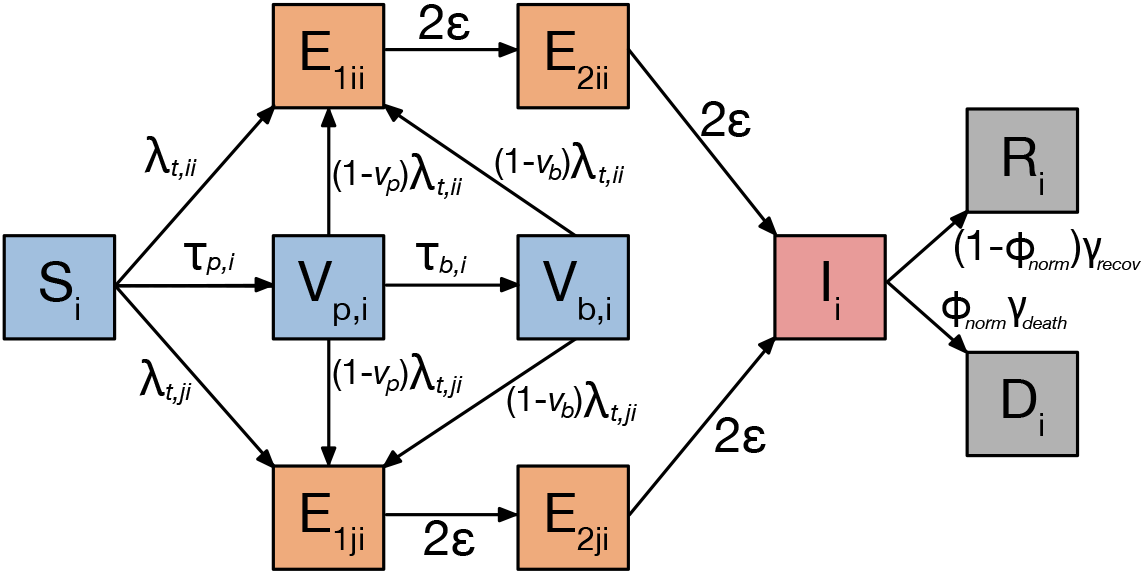
Schematic of the model structure. The population is stratified by occupation, so *i* and *j* are HCW (h) or community members (c). Individuals begin susceptible to infection (*S_i_*), and on infection they enter an exposed class (*E_i_*) split by the route of infection (*E_ii_, E_ji_*). There are 2 sequential *E* compartments so that the duration of the latent period is Erlang-distributed (see Methods). After the *E_2_* compartments, individuals enter the infectious compartment *(I)*, and then die *(D_i_*) or recover *(R_i_*). The force of infection, λ, depends on the route of transmission. When vaccination campaigns are implemented, susceptible individuals can enter the prime (*V_p,i_*) and boost (*V_b,i_*) compartments, and are then subject to a lower force of infection equal to 1-vaccine efficacy (*v_p_* or *v_b_*).

In the type 1 scenario, reactive strategies began on day 20 (April 27^th^), which is when health authorities were alerted to an outbreak of bloody diarrhoea in Kikwit [17]. This was earlier than detection of EVD, but we assumed that there would be improved surveillance and quicker EVD confirmation compared with 1995. For type 2 simulations, early transmission was slower. We therefore started reactive vaccination when the number of cases was similar to the type 1 outbreak on day 20, which was day 40 (median=53 cases in type 1, 38 in type 2). At 40 days, no simulation of the type 2 scenario had more than 100 cases. This is similar to the number of reported cases at commencement of vaccination in the recent outbreak in Nord Kivu (DRC)[28].

In ahead-of-time vaccination strategies of type 1 outbreaks the number of exposed and infected HCW at the start of the epidemic simulation were drawn from independent Poisson distributions with means from the joint posterior. For type 2 outbreaks, epidemics were seeded with 5 infected and 5 exposed community members (Supplementary Section S4.5).

### Sensitivity analysis

The type 2 scenario from published parameter estimates implies that HCW have lower onward transmission than community members. To explore this assumption, we conducted a sensitivity analysis where HCW transmission mirrors the transmission characteristics of community members, where both have moderate transmission and a later time of decrease in transmission (Supplementary Section S7).

## Results

### Classification of outbreaks into two types

Using key characteristics of the HCW and community transmission dynamics, we classified twelve localised EVD outbreaks into two broad types (Figure 2, Table 1, and Supplement S5). In both outbreak types, HCW were at high risk of infection, however, in type 1 outbreaks there was an early rapid increase in HCW incidence and in the cumulative proportion of HCW infected. Type 1 outbreaks also had a higher total proportion of HCW infected, shorter duration of HCW infections, and a smaller total outbreak size. In contrast, type 2 outbreaks exhibited a lower overall proportion of HCW infected, and a less obvious time period of high HCW incidence, with longer period of HCW infections. These outbreaks also showed a lower overall proportion of HCW infected, and larger total outbreak size.

The outbreak types are broad classifications, based on a combination of features of the dynamics of HCW infections. By classifying outbreaks in this way, we were able to determine the effect of HCW-targeted vaccination strategies under the range of observed transmission scenarios.

### Fit of model to Kikwit data

Our fitted model captured the dynamics of EVD in Kikwit (type 1 outbreak) for each route of transmission (Supplementary section S4). We found that initial transmission from HCW was high (median *R_hh_*=2.68, *R_hc_*=3.99) (Table 1 and Figure 4). In contrast, the within-community reproduction number was less than one, and therefore transmission was not sustainable. Although there was low *per capita* transmission from the community to HCW (median *R_ch_*=0.16), this represents a considerable risk to HCW: on average, each eight community cases infected one HCW. Overall, the net reproduction number at the start of the study period was 2.98 (2.11-4.36), with a major contribution from HCW, despite their low number. The timing and shape of the decrease in transmission depended on the occupation of cases (Figure 4 and Table 1). We inferred an early and rapid decrease in HCW-related transmission, however we found that within-community transmission decreased several weeks later. The net reproduction number fell below one after 30 (27-35) days (Figure 4c). Stochastic simulations of the type 1 scenario resulted in 288 (180-406) cases (observed value=284), and the final case was reported on day 115 (93-155) (Figure 5).

**Figure 4.**
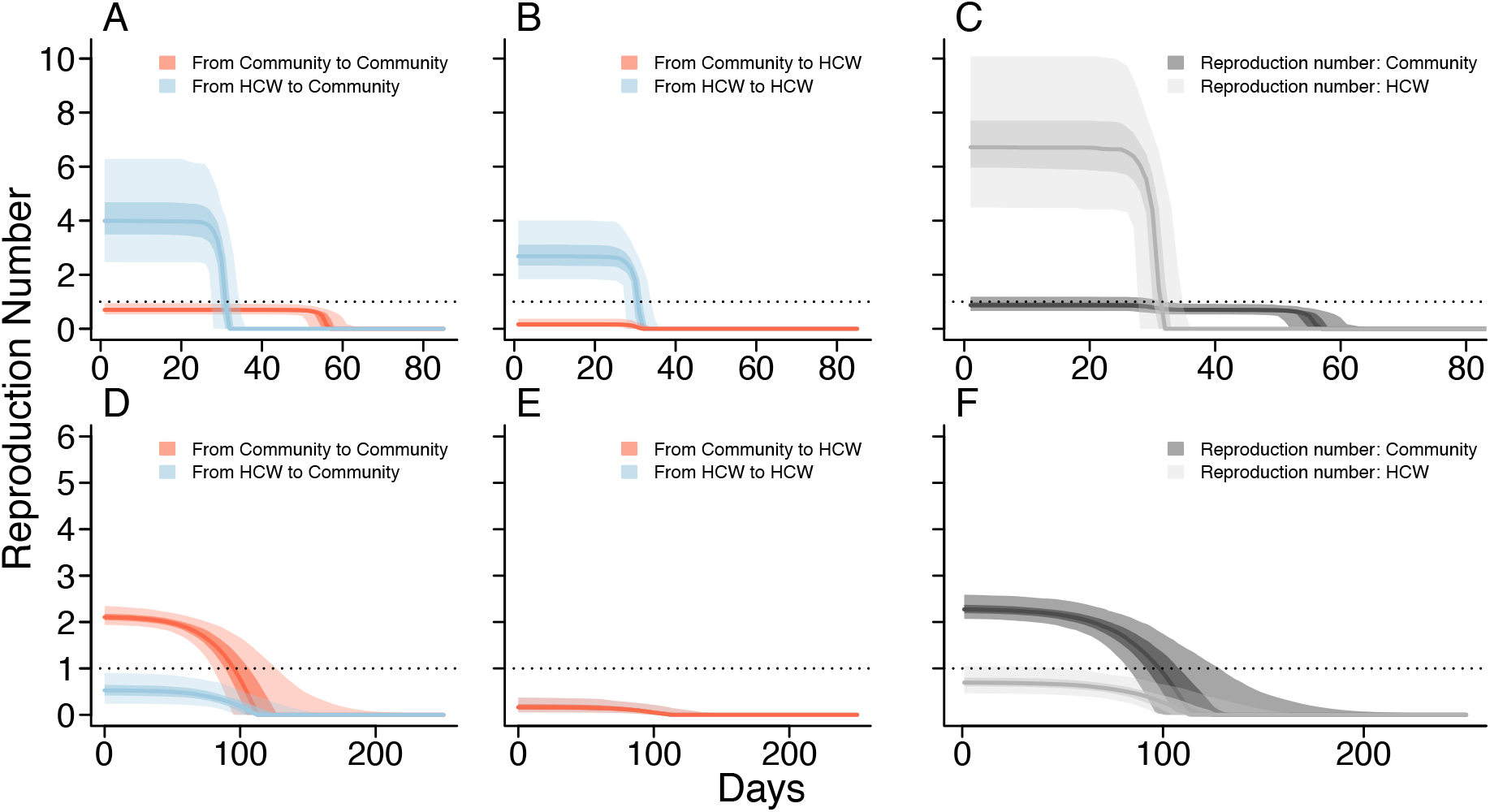
Reproduction number trajectories used in simulation for type 1 (upper) and type 2 (lower) outbreaks. A and D) HCW-to-community reproduction number (blue) and community-to-community (red) are given with mean and 50% and 95% CI. B and E) HCW-to-HCW reproduction number (blue) and community-to-HCW reproduction number (red) decrease at the same time, *T_h_* The horizontal line indicates *R* = 1. C and F) The overall reproduction number trajectories for Community members (dark grey) and HCW (light grey), with 50% and 95% CI.

**Figure 5.**
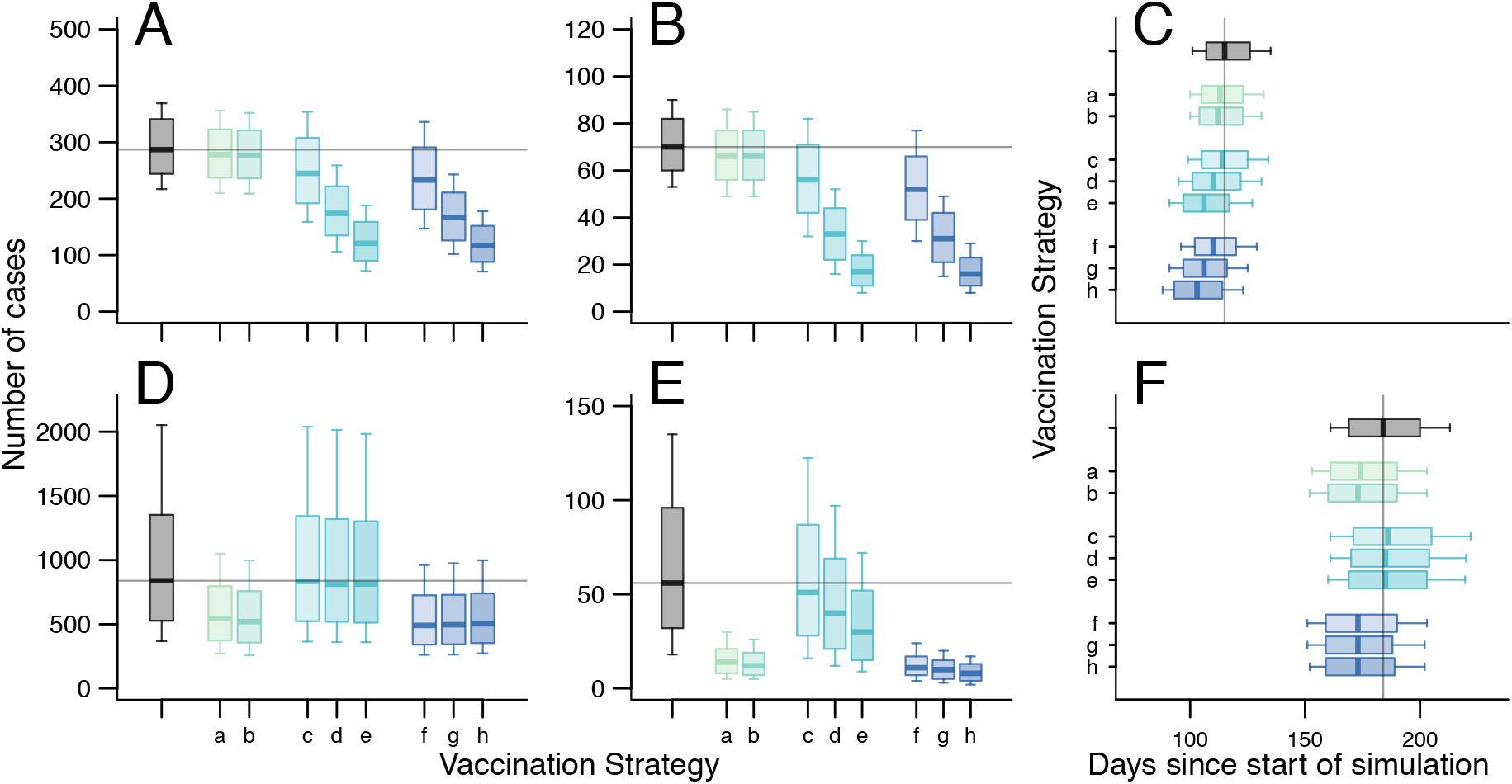
Effect of vaccination by vaccine strategy in type 1 (upper) and type 2 (lower) outbreaks. In type 1 outbreaks A) Number of cases in the entire population, B) number of cases in the 900 simulated HCW, and C) time to extinction; In type 2 outbreaks the: D) Number of cases in the entire population, E) number of cases in the 900 simulated HCW, and F) time to extinction. Boxplots show median value, rectangles mark 50% CI, and whiskers the 75% CI. Simulations without vaccination are shown in grey, and each colour represents a vaccination strategy (Table 3): reactive mass vaccination with (a) prime-boost vaccine or (b) single dose vaccine; ahead-of-time HCW vaccination only, with coverage in HCW of (c) 10%, (d) 30% or (e) 50%; ahead-of-time HCW vaccination plus reactive mass vaccination, with coverage in HCW of (f) 10%, (g) 30% or (h) 50%. We give the 75% CI due to high variation in the simulation sets, and 95% CI are given in the Supplement (S6.5) Note different y-axes.

### Comparison to type 2 scenario

Although it was not possible to fit the model directly to a type 2 outbreak, simulations of the type 2 scenario resulted in similar characteristics to those observed (Figure 2), matching patterns of cumulative proportion of HCW infected either through time or in total, as well as characteristics of number infected at each stage of the outbreak (Supplement S5).

As observed in the data, type 2 outbreaks were larger than in type 1 (839 cases (170-4473)) and the epidemics lasted longer (184 (149-229) days). In the baseline simulation, 70 (41-105) HCW were infected in type 1 outbreaks (observed value = 73), whereas 56 (5-263) were infected in the type 2 scenario (Figure 5).

### Ahead-of-time HCW-targeted vaccination strategies

These strategies had greater effect on total outbreak size in type 1 scenarios, whereas reactive and combined strategies had greater effect in type 2 scenarios (Figure 5). Impact of ahead-of-time HCW-targeted strategies depended on the coverage achieved: 50% coverage of HCW decreased the total number of cases in type 1 outbreaks (type 1: 121 (50-243), type 2: 813 (163-4245)). This strategy did not markedly shorten the outbreaks. This strategy did decrease the number of HCW infected in both type 1 and type 2 scenarios (Figure 5b and e). At lower coverage, there was less impact on total cases or duration, but there was a decrease in cases in HCW. The effect on simulated outbreak trajectories is shown in Supplementary Section S6.

### Reactive vaccination strategies

In type 1 outbreaks, this strategy was of limited benefit either to the entire population (Figure 5a) or to HCW (Figure 5b). In contrast, in type 2 simulations, vaccination substantially decreased the total cases, and there were 297 (7-2843) fewer cases in the entire population. Of those, 43 (3-226) fewer cases were HCW. Few type 2 simulations generated very large outbreaks with reactive mass vaccination (12% resulted in more than 1000 cases). In both type 1 and 2 scenarios, the single dose vaccine resulted in fewer cases than prime-boost, because of the delay until the boost dose, although the difference was small: 3 (−26-35) cases in type 1, and 19 (−109-235) in type 2.

### Combined vaccination strategies

Combined strategies decreased the number of cases and shortened the outbreak for all values of coverage. In the type 1 scenario, combined strategies did not decrease the number of HCW cases compared with ahead-of-time HCW vaccination, because reactive strategies began too late to protect additional HCW. In the type 2 scenario, for all values of HCW coverage, more than 80% of the simulations excluded the baseline median (839 cases), and time-to-extinction was reduced. In contrast to the type 1 scenario, the combination strategies provide extra protection to HCW directly, because reactive campaigns are assumed to prioritize HCW.

### Sensitivity analysis on HCW-related transmission in type 2 scenario

The overall size of outbreaks was smaller because of the lower total reproduction number (Supplementary Section S7). The general pattern of effect of different vaccination strategies was the same as the type 2 scenario parameterised to [21], which gives confidence in the generalisability of our findings.

## Discussion

We used as much information on HCW-related transmission as possible to classify EVD outbreaks into two broad types: the first, where the infection is catalysed by HCW and community transmission is low, was observed in Kikwit (1995) and some prefectures of Guinea and counties of Liberia during the large West African epidemic (2013-16); and the second, where there is high risk to HCW, but the epidemic is not amplified by their transmission, was observed in other areas of West Africa during the 2013-16 epidemic. We parameterised the type 1 scenario by fitting to an exemplar outbreak, and the type 2 scenario using published values for transmission between groups. This classification is not perfect, with some outbreaks exhibiting characteristics of both types, but allowed us to explore the range of observed infection dynamics relating to HCW infections and quantify the impact of HCW-targeted vaccine strategies in observed scenarios.

Using a mechanistic transmission model stratified by occupation and route of transmission, we found that in type 1 outbreaks, ahead-of-time HCW vaccination can have a large impact on the number of cases. Direct protection prevented infection of HCW, but also decreased their role in further spread. In these scenarios, where transmission is more dependent on the health care setting and perhaps, therefore more amenable to rapid decreases in transmission, there are limited additional benefits of reactive mass vaccination, both in number of cases averted, and the duration of the outbreak. Indeed, the model suggests that ahead-of-time vaccination of health care workers, even at modest coverage (30% immunised) is more effective than mass vaccination in response to outbreaks. Ahead-of-time HCW vaccination strategies require many fewer doses than mass vaccination strategies.

In type 2 scenarios, where within-community transmission is above the epidemic threshold, and there is no early decrease in HCW-related transmission, ahead-of-time vaccination of HCW could still provide individual protection to HCW and had a modest impact on overall transmission. However, reactive community vaccination (with or without ahead-of-time HCW vaccination) is more effective under these circumstances as this contributes to reducing the reproduction number below one.

In all modeled scenarios, ahead-of-time vaccination of HCW provided direct protection for HCW, and also decreased the number of cases in HCW due to indirect protection. In this analysis, we used the effective vaccine coverage, because we could not distinguish 30% protection of all HCW from 100% protection of 30% of the HCW. Further information on the likely protective effect of future vaccines would allow more specific examination of this distinction.

Data from Kikwit (1995) provide uniquely detailed information on likely source of infection and timing of symptoms. However, some data were missing, and the suggested routes of infection may not be correct. We did not consider transmission after death, or that some individuals (such as carers) may be more likely to be infected, which could affect estimates of transmission rate.

When generalising our findings to the current context, diagnosis and testing may now occur sooner than during the 1995 Kikwit outbreak. In addition, the rapid change in HCW-related transmission may partially have resulted from an increase in use of PPE. In the current context, HCW may have more rapid access to PPE, or have improved awareness of EVD, and therefore the change in transmission rate could be different. This would decrease the impact of HCW-targeted vaccination in type 1 scenarios, and therefore our findings may be on the upper end for ahead-of-time HCW vaccination.

We used data for all the outbreaks and sub-outbreaks within a larger epidemic that had information on the occupation of cases. It is possible that there is some misclassification of occupation, or that the definition of HCW changed from one outbreak to another, especially during the long West African epidemic. Incorrect classification could affect the reproduction numbers attributed to each group, although we do not have evidence of systematic misclassification. We conducted sensitivity analyses on the number of HCW and found that it did not affect the findings of vaccine impact.

Our model framework could not test other potential vaccination strategies, such as ring vaccination, because we did not track specific contacts that individuals make. We assumed that individuals mix randomly within occupation groups, with no heterogeneity within groups. Despite this limitation, the general conclusions are robust to the precise value of the number of HCW, and the transmission values we used.

Reactive HCW-targeted vaccination was used successfully during the 2018 DRC outbreaks [29], however, reactive vaccination strategies are logistically challenging. We incorporated delays into the model, but better information on the time from notification to vaccination would improve estimates of impact. These delays could also be affected by community response to vaccination, where resistance, and slowing of campaigns, could decrease vaccine impact.

Achieving ahead-of-time coverage may be challenging, due to high rates of turnover of health care staff, the large geographic area at risk of outbreaks, and because the duration of protection is currently unclear. However, strategies that target HCW in towns or cities nearby to an emerging outbreak are a potential way of achieving ahead-of-time coverage, as used in neighbouring areas of Uganda during the Nord Kivu outbreak [30], at the same time as implementing enhanced protective measures.

The categorisation proposed here is generated from a range of characteristics of EVD outbreaks. Not all EVD outbreaks are of these two types, since there are not always HCW infections [31]. In those cases, HCW vaccination would likely not improve outbreak control by decreasing transmission, although would provide direct protection to individuals at risk. However, we explored the impact of different assumptions about HCW-related transmission during sensitivity analysis, and found the general pattern of impact of each strategy to be similar.

Much of the data are drawn from local outbreaks during the larger West African epidemic, where a large number of interventions were occurring at different times and locations, and therefore care must be taken when extrapolating to new outbreak settings. Nevertheless, these local outbreaks exhibited different HCW-related dynamics, suggesting vaccine-led interventions could have varying impacts in different locations.

It would be challenging to assign outbreak classifications in real-time, because the division relies on some metrics available only after the outbreak has ended. Further work could develop a classification system more suited to use in real-time. However, we did not determine any adverse effects on number of cases or duration caused by any vaccine strategy, so the number of doses is the discriminant factor. Collecting and publishing more detailed information on occupation of cases and the route of transmission in future outbreaks would greatly improve our understanding of the epidemiology of EVD and the potential benefits from control measures targeted at different transmission routes.

Although we do not know whether the next outbreak will be type 1, type 2, or a mixed-type outbreak, ahead-of-time HCW vaccination decreased the number of cases seen in HCW in simulations of both outbreak types. Ahead-of-time HCW vaccination decreased the total number of cases to a small degree in type 2 outbreaks, but had a larger effect on type 1 outbreaks, by indirectly protecting non-vaccinated HCW and the community, even at modest levels of HCW coverage. Supplemental reactive community vaccination strategies may be required to control outbreaks when within-community transmission is intense, as seen in type 2 outbreaks. This analysis quantifies the impact of realistic and feasible vaccination strategies which may be implemented in a future EVD outbreak.

## Supporting information

## Authors’ contributions

AR^1^, RME, AC and WJE developed the analysis plan; JJMT, AR^2^ and KS collected and cleaned data from 1995 Kikwit outbreak, and 2013-2016 Guinea outbreak; AR^1^ implemented the analysis and ran the model, with contribution from AC; AR^1^ and RME interpreted the results, wrote the first draft and the supplemental material. AC, WJE, MB, AR^2^, JJMT, KS contributed to the manuscript. All authors approved the final version.

## Acknowledgements

We acknowledge Sebastian Funk for technical support.

## Ethics approval

This study is a secondary analysis of previously collected data.

## References

1. World Health Organization. Ebola Situation Report March 30, 2016. 2016; 1–16.

2. Grinnell M, Dixon MG, Patton M, et al. Ebola Virus Disease in Health Care Workers - Guinea, 2014. MMWR 2015; 64:1083–7.

3. Matanock A, Arwady MA, Ayscue P, et al. Ebola Virus Disease Cases Among Health Care Workers Not Working in Ebola Treatment Units — Liberia, June – August, 2014. Morb Mortal Wkly Rep 2014; 63:1–5.

4. Olu O, Kargbo B, Kamara S, et al. Epidemiology of Ebola virus disease transmission among health care workers in Sierra Leone, May to December 2014: a retrospective descriptive study. BMC Infect Dis 2015; 15:416.

5. Kilmarx PH, Clarke KR, Dietz PM, et al. Ebola virus disease in health care workers--Sierra Leone, 2014. MMWR Morb Mortal Wkly Rep 2014; 63:1168–71.

6. Khan AAS, Tshioko FKF, Heymann DLD, et al. The Reemergence of Ebola Hemorrhagic Fever, Democratic Republic of the Congo, 1995. J Infect Dis 1999; 179:S76–86.

7. World Health Organization. Health worker Ebola infections in Guinea, Liberia and Sierra Leone-A preliminary report. 2015; 1–16.

8. Kucharski AJ, Camacho A, Flasche S, Glover RE, Edmunds WJ, Funk S. Measuring the impact of Ebola control measures in Sierra Leone. Proc Natl Acad Sci U S A 2015; 112:14366–71. Available at: http://www.pnas.org/content/112/46/14366.abstract?sid=719d46c3-50db-4bbc-a38b-5cd4f3e243c9. Accessed 21 February 2016.

9. Camacho A, Kucharski AJ, Funk S, Breman J, Piot P, Edmunds WJ. Potential for large outbreaks of Ebola virus disease. Epidemics 2014; Available at: http://www.sciencedirect.com/science/article/pii/S1755436514000528. Accessed 7 October 2014.

10. Legrand J, Grais RF, Boelle P-Y, Valleron AJ, Flahault A. Understanding the dynamics of Ebola epidemics. Epidemiol Infect 2007; 135:610. Available at: http://www.journals.cambridge.org/abstract_S0950268806007217.

11. Kucharski AJ, Camacho A, Checchi F, et al. Evaluation of the benefits and risks of introducing Ebola community care centers, Sierra Leone. Emerg Infect Dis 2015; 21:393–9.

12. WHO Ebola Response Team. Ebola Virus Disease in West Africa — The First 9 Months of the Epidemic and Forward Projections. N Engl J Med 2014; 371:1481–1495. Available at: http://www.nejm.org/doi/abs/10.1056/NEJMoa1411100.

13. Merler S, Ajelli M, Fumanelli L, et al. Containing Ebola at the Source with Ring Vaccination. PLoS Negl Trop Dis 2016; 10:e0005093.

14. Kucharski AJ, Eggo RM, Watson CH, Camacho A, Funk S, Edmunds WJ. Effectiveness of Ring Vaccination as Control Strategy for Ebola Virus Disease. Emerg Infect Dis 2016; 22:105–8. Available at: http://www.pubmedcentral.nih.gov/articlerender.fcgi?artid=4696719&tool=pmcentrez&rendertype=abstract.Accessed 19 February 2016.

15. World Health Organization. Fact sheet no. 103 Ebola virus disease. 2014.

16. Rosello A, Mossoko M, Flasche S, et al. Ebola virus disease in the Democratic Republic of the Congo, 1976-2014. Elife 2015; 4:e09015. Available at: http://elifesciences.org/content/4/e09015.abstract.Accessed 3 November 2015.

17. Muyembe-Tamfum JJ, Kipasa M, Kiyungu C, Colebunders R. Ebola outbreak in Kikwit, Democratic Republic of the Congo: discovery and control measures. J Infect Dis 1999; 179 Suppl:S259–S262.

18. King AA, Domenech de Celles M, Magpantay FMG, Rohani P. Avoidable errors in the modelling of outbreaks of emerging pathogens, with special reference to Ebola. Proc R Soc B Biol Sci 2015; 282:20150347–20150347. Available at: http://rspb.royalsocietypublishing.org/content/282/1806/20150347.Accessed 1 April 2015.

19. Roberts GO, Rosenthal JS. Examples of adaptive MCMC. J Comput Graph Stat 2009; 18:349–367.

20. Andrieu C, De Freitas N, Doucet A, Jordan MI. An introduction to MCMC for machine learning. Mach Learn 2003; 50:5–43.

21. Faye O, Boëlle P-Y, Heleze E, et al. Chains of transmission and control of Ebola virus disease in Conakry, Guinea, in 2014: an observational study. Lancet Infect Dis 2015; 15:320–326. Available at: http://www.ncbi.nlm.nih.gov/pubmed/25619149.Accessed 20 February 2018.

22. Camacho A, Kucharski A, Aki-Sawyerr Y, et al. Temporal Changes in Ebola Transmission in Sierra Leone and Implications for Control Requirements: a Real-time Modelling Study. PLoS Curr 2015; 7. Available at: http://www.pubmedcentral.nih.gov/articlerender.fcgi?artid=4339317&tool=pmcentrez&rendertype=abstract. Accessed 6 July 2015.

23. Funk S, Ciglenecki I, Tiffany A, et al. The impact of control strategies and behavioural changes on the elimination of ebola from lofa county, Liberia. Philos Trans R Soc B Biol Sci 2017; 372.

24. Ajelli M, Parlamento S, Bome D, et al. The 2014 Ebola virus disease outbreak in Pujehun, Sierra Leone: epidemiology and impact of interventions. BMC Med 2015; 13:281.

25. Santermans E, Robesyn E, Ganyani T, et al. Spatiotemporal evolution of Ebola virus disease at sub-national level during the 2014 West Africa epidemic: Model scrutiny and data meagreness. PLoS One 2016;

26. Shoman H, Karafillakis E, Rawaf S. The link between the West African Ebola outbreak and health systems in Guinea, Liberia and Sierra Leone: a systematic review. Global Health 2017; 13:1.

27. Petit D, Sondorp E, Mayhew S, Roura M, Roberts B. Implementing a Basic Package of Health Services in post-conflict Liberia: Perceptions of key stakeholders. Soc Sci Med 2013; 78:42–49.

28. WHO. Ebola vaccination begins in North Kivu. 2018. Available at: http://www.who.int/news-room/detail/08-08-2018-ebola-vaccination-begins-in-north-kivu. Accessed 29 August 2018.

29. WHO. WHO supports Ebola vaccination of high risk populations in the Democratic Republic of the Congo. 2018. Available at: http://www.who.int/news-room/detail/21-05-2018-who-supports-ebola-vaccination-of-high-risk-populations-in-the-democratic-republic-of-the-congo. Accessed 27 July 2018.

30. Health UM of. UGANDA VACCINATES FRONTLINE HEALTHWORKERS AGAINST EBOLA. 2018. Available at: https://health.go.ug/download/file/fid/2035.

31. World Health Organization. Ebola Virus Disease Situation Report in DRC: 8 June 2017. 2017;

