## Supplementary material for "Effect of vaccinating health care workers to control Ebola virus disease: a modelling analysis of outbreak data"

**S7 Sensitivity analysis on HCW transmission in type 2 scenario** **22**

**Bibliography** **25**

### S1 Data from West African epidemic

This section examines data from the 2013-16 West African EVD outbreak to clarify the modes of transmission between HCW and community members.

#### S1.1 Guinea

We stratified data from Guinea by prefecture and compared the incidence among HCW and community. These data included the date of onset of symptoms, the location of the case (at a prefecture and sub-prefecture level), and their occupation (HCW or community member) and were drawn from the Guinea Ministry of Health database. HCW were defined according to the WHO definition, including not only clinical staff, but all those who work in health services (i.e. drivers, cleaners, burial teams, and community-based workers) [1].

The daily incidence of EVD stratified by prefecture highlights that several geographically-distinct outbreaks affected the country (Figure S1). The timeseries of these outbreaks show different transmission dynamics. In the urbanised capital Conakry, infection of HCW occurred throughout the long outbreak (Figure S1A). In Kissidougou we observed a shorted epidemic, with HCW affected solely at the start of the outbreaks. Whereas in Macenta, Coyah, and Guéckédou prefectures, transmission almost exclusively occurred among community members (14 HCW recorded out of 693 cases in Macenta prefecture, 19 out of 217 in Coyah and 11 out of 310 in Guéckédou). In Dubreka and Kerouane(Figure S1D and E) we suspect multiple re introductions occurred at a sub prefecture level, causing several spikes in the number of cases. Therefore Kerouane and Dubreka are not included in the analysis.

We classified the time series of EVD onset in Conakry, Coyah, Gueckedou, Kissidougou, N'Zérékoré and Macenta according to the dynamics of HCW infection in each setting (Figure S2). We observe all Guinean outbreaks but Kissidougou follow a similar pattern, i.e HCW transmission constantly observed throughout the outbreaks, which last for more than 100 days.

The time series in Conakry, Macenta and Gueckedou districts show a long-lasting transmission in urban settings, the incidence of EVD among HCW is higher than in the rest of the community, and occur throughout the outbreak. This dynamic is different than seen in Kikwit, which is a shorter outbreak with rapid increase in HCW infections.

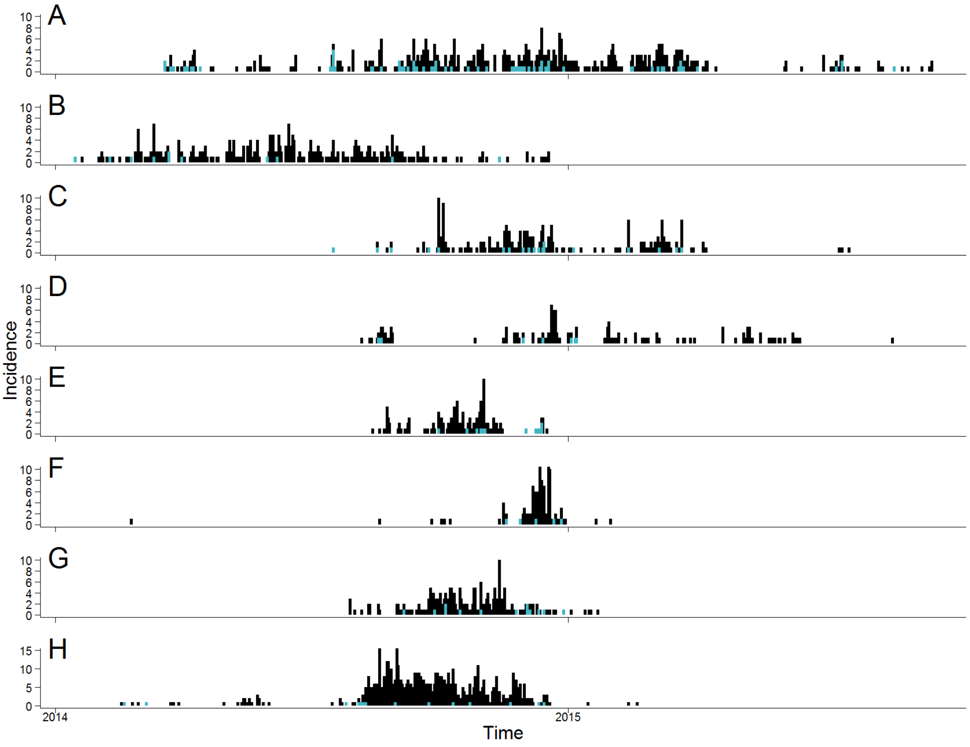

Figure S1: Daily incidence time series of EVD onset during the 2013-2016 outbreak in Guinea, split by prefecture : A) Conakry B) Guéckédou C) Coyah D) Dubréka E) Kerouane F) Kissidougou G) N'Zérékoré H) Macenta ; the cases are stratified by occupation (HCW : blue vertical line ; community : black vertical lines).

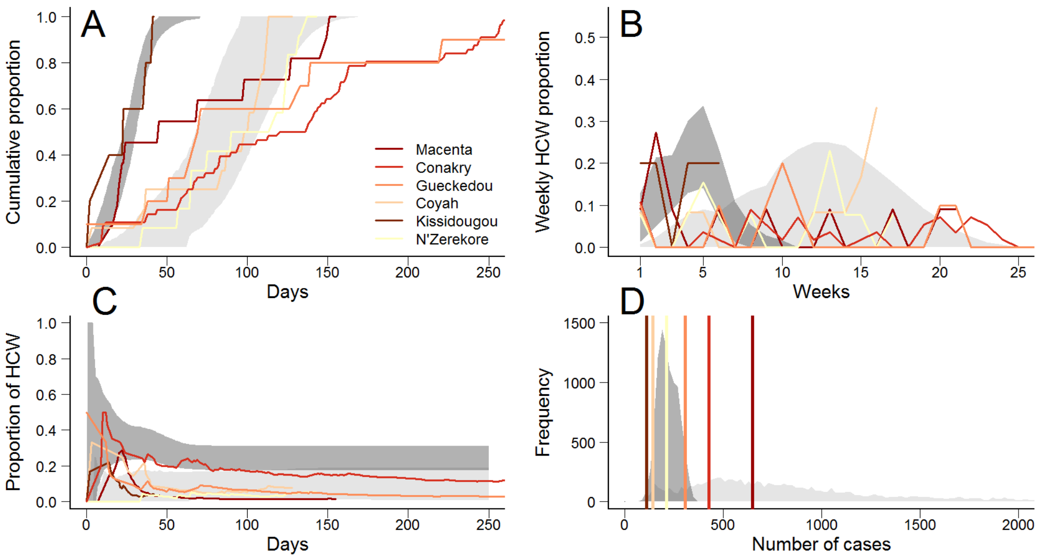

Figure S2: Classification of EVD outbreaks into two types according to their HCW-related dynamics. Dark grey marks the 95% CI in type 1 outbreak simulations, light grey areas correspond to 95% CI for type 2 outbreak simulations. A) Cumulative proportion of HCW among the total number of HCW infected through time. B) Weekly proportion of HCW among the total number of HCW infected in EVD outbreaks through time. C) Proportion of HCW infected through time. D) Distribution of the number of community cases, where vertical lines represent the number of community cases observed in each outbreak.

#### S1.2 Liberia

We calculated the daily cumulative number of cases from the 2013-2016 outbreak in counties of Liberia stratified by HCW and community (Figure S3). These data were drawn from publicly available sources of data, and were compiled into usable format and provided online [2]. In Bong County and Margibi County, the number of HCW cases rapidly increased in the early phase of the epidemic and then stabilised, suggesting a reduction in risk to HCW, while further cases occurred only in the community. This pattern is not discernible when looking at a national level, since EVD occurred in different regions at different times, thus masking the county-specific patterns. In Nimba County, the number of HCW involved is less than 10 and still increasing when this dataset ends, it is therefore not included in the rest of the analysis. HCW were defined according to the WHO definition, including not only clinical staff, but all those who work in health services (i.e. drivers, cleaners, burial teams, and community-based workers) [1].

The figure S4 highlights the similarity between the dynamics observed in Bong, Margibi and Bomi counties and in Kikwit. These time series highlight different dynamics among HCW and the rest of the population, and are reminiscent of the pattern observed in Kikwit, that is: early propagation among HCW, followed by a rapid decrease in transmission. On the other hand, a far slower increase was observed in Montserrado county, which is closer to type 2 outbreaks. Furthermore, far more cases were notified in Montserrado than in the other Liberian counties, highlighting another difference between type 1 and type 2 EVD outbreaks.

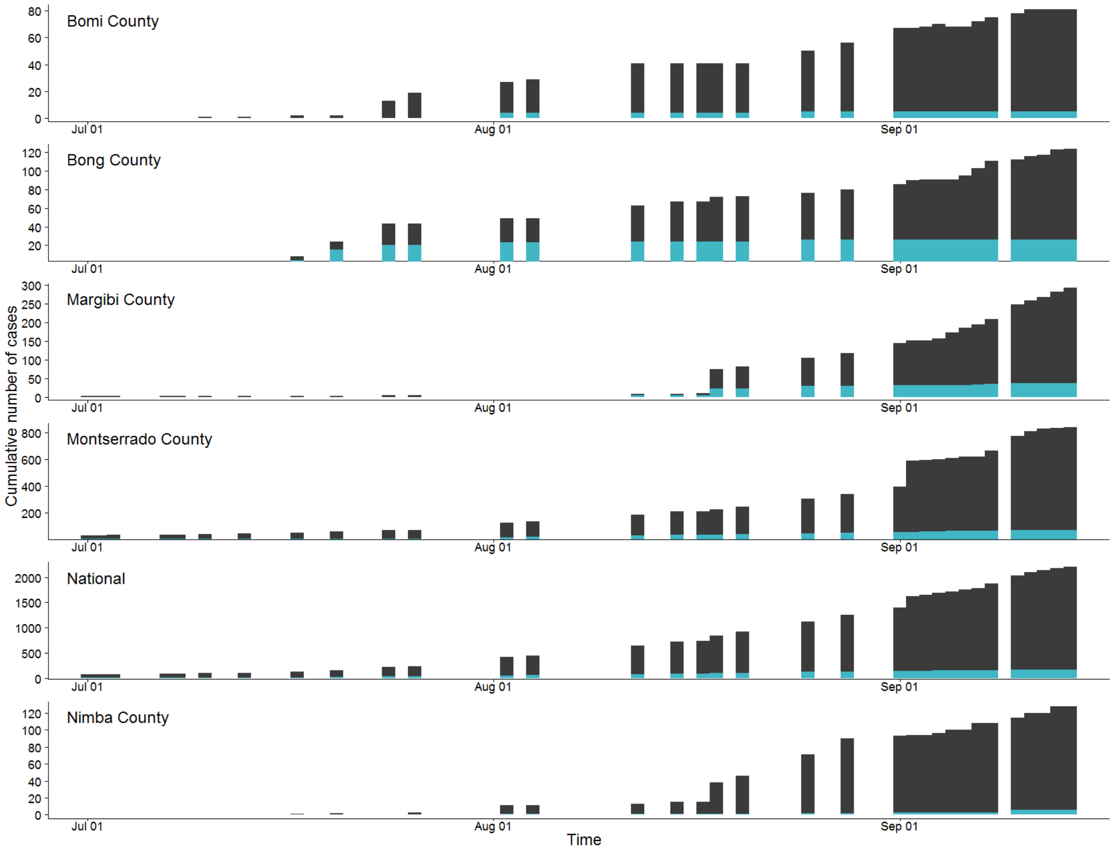

Figure S3: Cumulative number of confirmed and probable cases during the 2013-2016 Ebola outbreak in Liberia : A) Bomi County B) Bong County C) Margibi County D) Montserrado County E) National level F) Nimba County ; the cases are stratified by occupation (HCW (blue), community (black)).

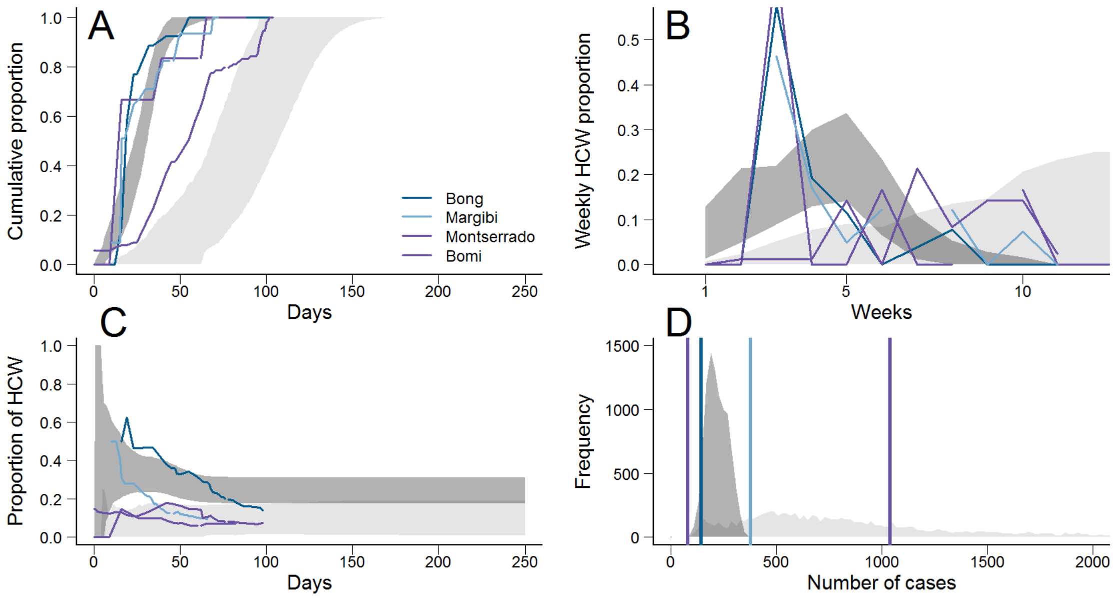

Figure S4: Classification of EVD outbreaks into two types according to their HCW-related dynamics. Dark grey marks the 95% CI in type 1 outbreak simulations, light grey areas correspond to 95% CI for type 2 outbreak simulations. A) Cumulative proportion of HCW among the total number of HCW infected through time. B) Weekly proportion of HCW among the total number of HCW infected in EVD outbreaks through time. C) Proportion of HCW infected through time. D) Distribution of the number of community cases, where vertical lines represent the number of community cases observed in each outbreak.

#### S1.3 Sierra Leone

We calculated the weekly onset of EVD by occupation is shown in Kenema District, Sierra Leone (Figure S5, [3]). HCW were defined as anyone who worked in a healthcare facility or engaged in healing practices [3]. We observe that in the early stages of the outbreak there was a high proportion of HCW cases, but the cumulative number of reported HCW cases flattened two months after the start of the outbreak, whereas transmission in the community was still occurring (Figure S6). The proportion of HCW infected therefore decreases over time. The Kenema setting was unusual as strikes among HCW occurred after 6 weeks [3], possibly changing the dynamics of transmission. HCW dynamics in Kenema seem to differ from Kikwit setting.

Various features of the Kenema EVD outbreak are displayed in Figure S6, which highlights HCW infections lasted for more than 100 days. More than 500 cases were reported during Kenema outbreak. Type 2 outbreaks tend to be larger in size than type 1.

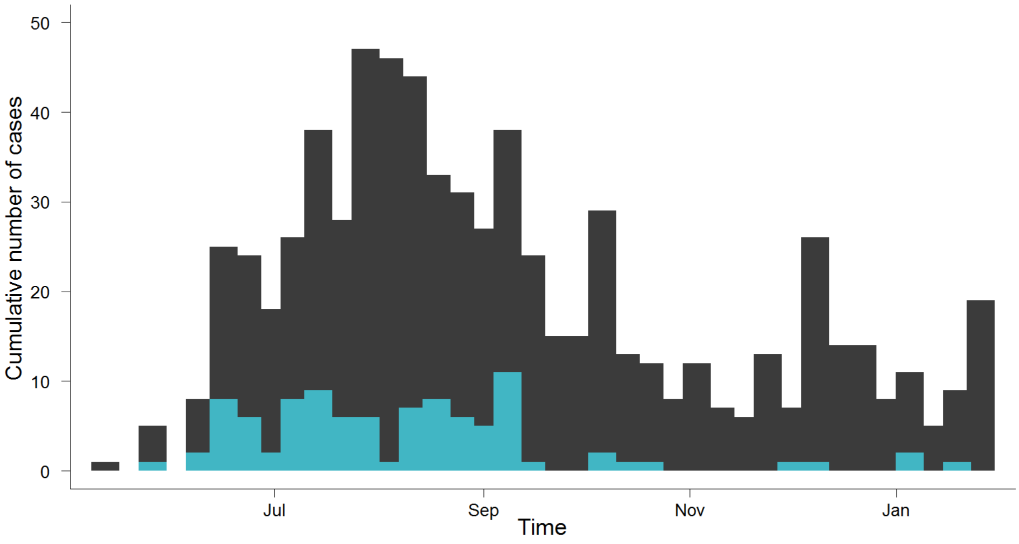

Figure S5: Weekly incidence time series of EVD onset during the 2013-2016 outbreak in Kenema in Sierra Leone.

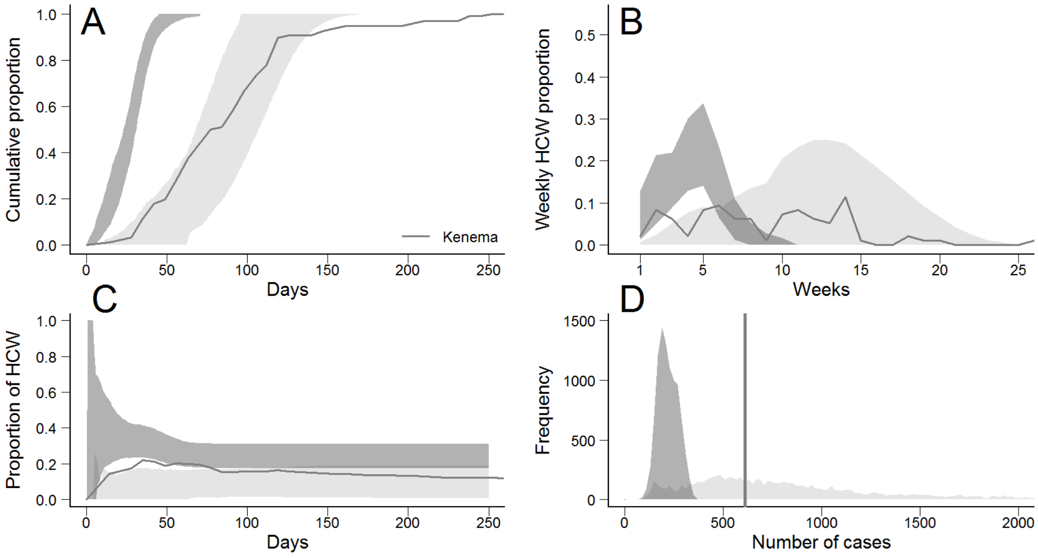

Figure S6: Classification of EVD outbreaks into two types according to their HCW-related dynamics. Dark grey marks the 95% CI in type 1 outbreak simulations, light grey areas correspond to 95% CI for type 2 outbreak simulations. A) Cumulative proportion of HCW among the total number of HCW infected through time. B) Weekly proportion of HCW among the total number of HCW infected in EVD outbreaks through time. C) Proportion of HCW infected through time. D) Distribution of the number of community cases, where vertical lines represent the number of community cases observed in each outbreak.

### S2 Further description of Kikwit data

An individual admitted to Kikwit General Hospital on April $7^{th}$ 1995, underwent surgery on April $10^{th}$, and was retrospectively confirmed as an EVD case. The first clinical diagnosis of haemorrhagic fever was made on May $2^{nd}$, leading to local awareness campaigns and control measures. EVD was confirmed on May $8^{th}$. On May $10^{th}$, WHO mobilized international assistance and further control measures were taken, such as the arrival of new protective equipment for HCWs, the opening of isolation wards, population awareness of the disease, and the communication campaign in the population [4]. The last case died on July $16^{th}$ 1995. In total 317 cases were reported, with 248 deaths (Case Fatality Ratio (CFR)=78%). Our study begins on April $7^{th}$, after which there were 284 cases, of which 73 are in HCW. There were several missing dates in onset and death (Table S1). HCW were defined as the workers on the personnel list of the hospitals [5].

| Total dataset | | | | | |
|---|---|---|---|---|---|
| Occupation | Total Number of cases | Date of onset given | Number of death | Date of death given | Index case given |
| HCW | 76 | 74 | 59 | 59 | 50 |
| Community | 241 | 213 | 189 | 187 | 173 |
| After arrival of index case in Kikwit General Hospital (7 April) | | | | | |
| Occupation | Total number of cases | Date of onset given | Number of deaths | Date of death given | Index case given |
| HCW | 73 | 67 | 58 | 58 | 48 |
| Community | 211 | 192 | 163 | 162 | 143 |

Table S1: Summary of the data. The values in the total dataset were used to compute epidemiological features (reporting fraction, case fatality ratio, etc), the data we used to fit the model was cases occurring after 7 April.

The number of cases in each stratification of infector-infectee pair, for onset or death is: the onset date of community cases with a community infector (within-community transmission, 88 cases), the onset date of HCW cases with a community infector (community-to-HCW transmission, 6 cases), the onset date of community cases with an HCW infector (HCW-to-community transmission, 55 cases), the onset date of HCW cases with an HCW infector (within-HCW transmission, 42 cases), the onset date of community cases with an unknown infector (unknown-to-community transmission, 49 cases), the onset date of HCW cases with an unknown infector (unknown-to-HCW transmission, 19 cases), the reported deaths among the community (162 deaths), and the reported deaths among HCW (58 deaths).
Kikwit had an approximate population of 200,000 in 1995 [6], and 429 HCW were employed at Kikwit General Hospital [5], which was the largest of its two hospitals. By 2003, 1047 HCW were employed in these two hospitals [7]. There was no information on the total number of HCW in other health care facilities surrounding Kikwit, so we estimated there were 900 HCW in total in 1995. Our findings were not sensitive to this value (Supplementary Section S4.3).

#### S2.1 Comparison of cases by occupation

We performed analysis to determine if duration of infection, CFR, and fraction of missing dates were the same in the two occupation groups (HCW and community) (Table S2). Using a binomial test to compare proportions, and a two sided student t-test for durations, we found the values in community cases and HCW were not significantly different for these variables. Therefore we used the same value for CFR, duration of infection and report rates for both populations (Table S2). We estimated the mean times from onset to recovery or death from the data. In Figure S7 we display the distributions of the durations between the onset of symptoms and the date of death / recovery in the two populations. The distributions were not significantly different in each population.

| Parameter | Pop C | Pop H | $p$-value |
|---|---|---|---|
| Duration onset to death (days) | 9.42 | 9.77 | 0.73 |
| Duration onset to recovery (days) | 17.55 | 20.37 | 0.50 |
| Case-fatality Ratio | 0.79 | 0.77 | 0.78 |
| Proportion of onset dates reported | 0.89 | 0.97 | 0.66 |
| Proportion of death dates reported | 0.98 | 1 | 1 |
| Proportion of recovery dates reported | 0.56 | 0.70 | 0.33 |

Table S2: Comparison of epidemiological features depending on occupation. We used a binomial test to compare the proportion of reported dates in the two populations, and a two-sided student t-test to compare the distribution of durations.

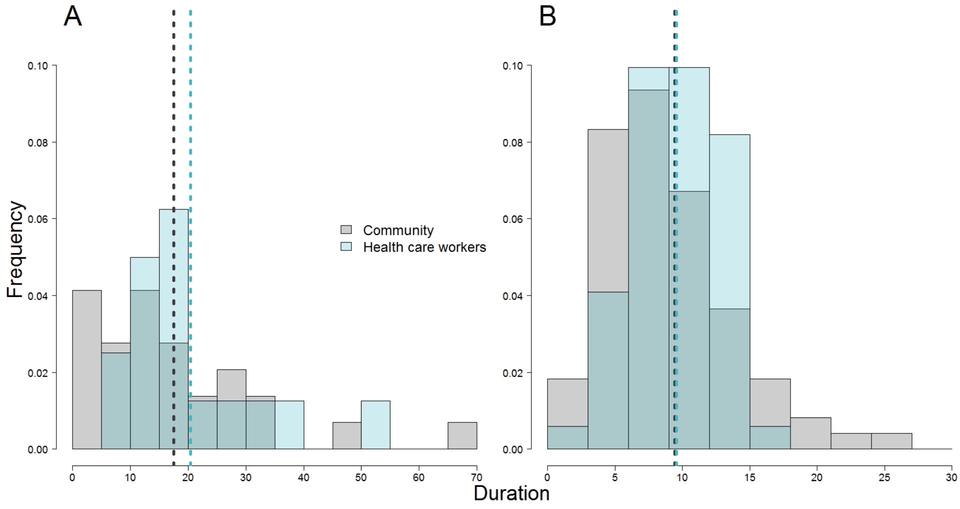

Figure S7: A. Distribution of the duration between the date of onset of symptoms and the date of recovery in Community and in HCW. B. Distribution of the duration between the date of onset of symptoms and the date of death in Community and in HCW. The vertical dotted lines represent the mean value for each population.

### S3 Modelling framework and inference

#### S3.1 Time-dependent change in transmission rate

To account for the effect of control measures such as the arrival of protective equipment for HCWs, the opening of isolation wards, and population awareness of EVD, we used flexible time-dependent decreasing sigmoid functions for the transmissibility parameters, $\beta_{t,ij}$:

$$\beta_{t,ij} = \beta_{t_o,ij}(1 - \frac{\delta_k}{1 + e^{-\alpha_k(t-T_k)}})$$

where $\delta$ is the reduction in the transmission rate, $\alpha$ is the shape of the change, $t$ is time in days, $T$ is the midpoint of the time of change for each occupation group, and $k = h$ if either $i$ or $j$ is $h$, and $k = c$ otherwise. We estimated these parameters of the sigmoid function.

#### S3.2 Calculation of the Reproduction number

We use the next generation matrix (NGM) to calculate the reproduction number, where $M$ is the mean duration of infectiousness.

$$M = \phi\gamma_{death}^{-1} + (1 - \phi)\gamma_{recov}^{-1}$$

$$NGM(t) = \begin{pmatrix} R_{cc}(t) & R_{ch}(t) \\ R_{hc}(t) & R_{hh}(t) \end{pmatrix} = \begin{pmatrix} \beta_{cc}(t)M * N_c/N & \beta_{ch}(t)M * N_h/N \\ \beta_{hc}(t)M * N_c/N & \beta_{hh}(t)M * N_h/N \end{pmatrix}$$

The basic reproduction number $R_0$ is the largest eigenvalue of the next generation matrix at $t = T_0$ [8]. The outcome of the Infected class is split into two compartments, so we defined a normalized Case Fatality Ratio $\phi_{norm}$ so that the overall CFR is $\phi$:

$$\phi_{norm} = \frac{\phi\gamma_{recov}}{\phi\gamma_{recov} + (1 - \phi)\gamma_{death}}$$

#### S3.3 Model equations

The equations of the full model are as follows (for brevity, the time dependencies of state variables are omitted):

$$\frac{dS_c}{dt} = -(\beta_{cc}(t)I_c + \beta_{hc}(t)I_h)\frac{S_c}{N} - \tau_{pc} \qquad \frac{dS_h}{dt} = -(\beta_{ch}(t)I_c + \beta_{hh}(t)I_h)\frac{S_h}{N} - \tau_{ph}$$

$$\frac{dV_{pc}}{dt} = \tau_{pc} - (1-\nu_p)(\beta_{cc}(t)I_c + \beta_{hc}(t)I_h)\frac{V_{pc}}{N} - \tau_{bc} \qquad \frac{dV_{ph}}{dt} = \tau_{ph} - (1-\nu_p)(\beta_{ch}(t)I_c + \beta_{hh}(t)I_h)\frac{V_{ph}}{N} - \tau_{bh}$$

$$\frac{dV_{bc}}{dt} = \tau_{bc} - (1-\nu_b)(\beta_{cc}(t)I_c + \beta_{hc}(t)I_h)\frac{V_{bc}}{N} \qquad \frac{dV_{bh}}{dt} = \tau_{bh} - (1-\nu_b)(\beta_{ch}(t)I_c + \beta_{hh}(t)I_h)\frac{V_{bh}}{N}$$

$$\frac{dE_{1cc}}{dt} = \beta_{cc}(t)I_c\frac{S_c + (1-\nu_p)V_{pc} + (1-\nu_b)V_{bc}}{N} - 2\epsilon E_{1cc} \qquad \frac{dE_{1ch}}{dt} = \beta_{ch}(t)I_c\frac{S_h + (1-\nu_p)V_{ph} + (1-\nu_b)V_{bh}}{N} - 2\epsilon E_{1ch}$$

$$\frac{dE_{2cc}}{dt} = 2\epsilon E_{1cc} - 2\epsilon E_{2cc} \qquad \frac{dE_{2ch}}{dt} = 2\epsilon E_{1ch} - 2\epsilon E_{2ch}$$

$$\frac{dE_{1hc}}{dt} = \beta_{hc}(t)I_h\frac{S_c + (1-\nu_p)V_{pc} + (1-\nu_b)V_{bc}}{N} - 2\epsilon E_{1hc} \qquad \frac{dE_{1hh}}{dt} = \beta_{hh}(t)I_h\frac{S_h + (1-\nu_p)V_{ph} + (1-\nu_b)V_{bh}}{N} - 2\epsilon E_{1hh}$$

$$\frac{dE_{2hc}}{dt} = 2\epsilon E_{1hc} - 2\epsilon E_{2hc} \qquad \frac{dE_{2hh}}{dt} = 2\epsilon E_{1hh} - 2\epsilon E_{2hh}$$

$$\frac{dI_c}{dt} = 2(\epsilon E_{2cc} + \epsilon E_{2hc}) - (\gamma_{death}\phi_{norm} + \gamma_{recov}(1-\phi_{norm}))I_c \qquad \frac{dI_h}{dt} = 2(\epsilon E_{2ch} + \epsilon E_{2hh}) - (\gamma_{death}\phi_{norm} + \gamma_{recov}(1-\phi_{norm}))I_h$$

$$\frac{dD_c}{dt} = \gamma_{death}\phi_{norm}I_c \qquad \frac{dD_h}{dt} = \gamma_{death}\phi_{norm}I_h$$

$$\frac{dR_c}{dt} = \gamma_{recov}(1-\phi_{norm})I_c \qquad \frac{dR_h}{dt} = \gamma_{recov}(1-\phi_{norm})I_h$$

In order to account for the week before vaccine protection becomes effective, $\tau_{pc} = 0$ and $\tau_{ph} = 0$ while $T < T_v + 7$, with mass vaccination campaign starting on $T = T_v$. To compare the model output with observed data, we calculated the incidences corresponding to the eight time series on each day. $k_c(t)$ and $k_h(t)$ represent the proportion of new cases (from Community and HCW respectively) with index case given on day $t$. Hence the likelihood of the daily unassigned infections ($\Delta I_{na-c}(t)$ and $\Delta I_{na-c}(t)$) was computed without taking into account the index case occupation, whereas with $\Delta I_{i-j}(t)$ we used the number of exposed individuals from this route of transmission only (compartments $E_{2ij}$). The mean number of reported cases from community-to-community per day t is $\rho_{onset} * \Delta I_{c-c}(t)$ and the variance is $\rho_{onset} * \Delta I_{c-c}(t) + (\rho_{onset} * \Delta I_{c-c}(t))^2\psi_{onset}$.

$$\Delta I_{c-c}(t) = \int_t^{t+1} 2\epsilon E_{2cc}k_c(t)\,\mathrm{d}t \qquad \Delta I_{c-h}(t) = \int_t^{t+1} 2\epsilon E_{2ch}k_h(t)\,\mathrm{d}t$$

$$\Delta I_{h-c}(t) = \int_t^{t+1} 2\epsilon E_{2hc}k_c(t)\,\mathrm{d}t \qquad \Delta I_{h-h}(t) = \int_t^{t+1} 2\epsilon E_{2hh}k_h(t)\,\mathrm{d}t$$

$$\Delta I_{na-c}(t) = \int_t^{t+1} 2\epsilon(E_{2cc} + E_{2hc})(1-k_c(t))\,\mathrm{d}t \qquad \Delta I_{na-h}(t) = \int_t^{t+1} 2\epsilon(E_{2ch} + E_{2hh})(1-k_h(t))\,\mathrm{d}t$$

$$\Delta D_c(t) = \int_t^{t+1} \gamma_{death}\phi_{norm}I_c\,\mathrm{d}t \qquad \Delta D_h(t) = \int_t^{t+1} \gamma_{death}\phi_{norm}I_H\,\mathrm{d}t$$

### S4 Model output and sensitivity analysis

#### S4.1 Fit to Kikwit data

The eight timeseries used for the fit are displayed in Figure S8. These data were used to fit the model.

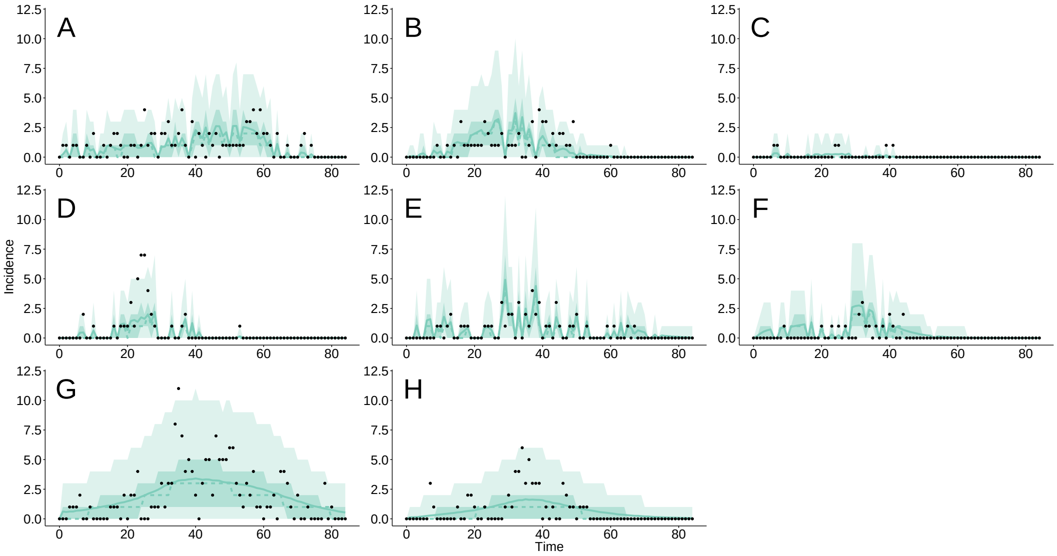

Figure S8: Comparison of our fitted model and observed daily incidence time series (black dots) reconstructed from the line list of EVD cases in Kikwit in 1995. A) Daily onset, from community-to-community; B) Daily onset, from HCW-to-community; C) Daily onset, from community-to-HCW ; D) Daily onset, from HCW-to-HCW ; E) Daily onset in the community, without information about index case ; C) Daily onset among HCW, without information about index case ; G) Daily deaths in the community ; H) Daily deaths among HCW. The mean and median fits are represented by solid and dashed lines respectively. The dark and light shaded areas correspond to the 50% and 95% credible intervals respectively.

The aggregated time series of date of onset and date of death in the entire population show very good agreement with observed values (Figure S9).

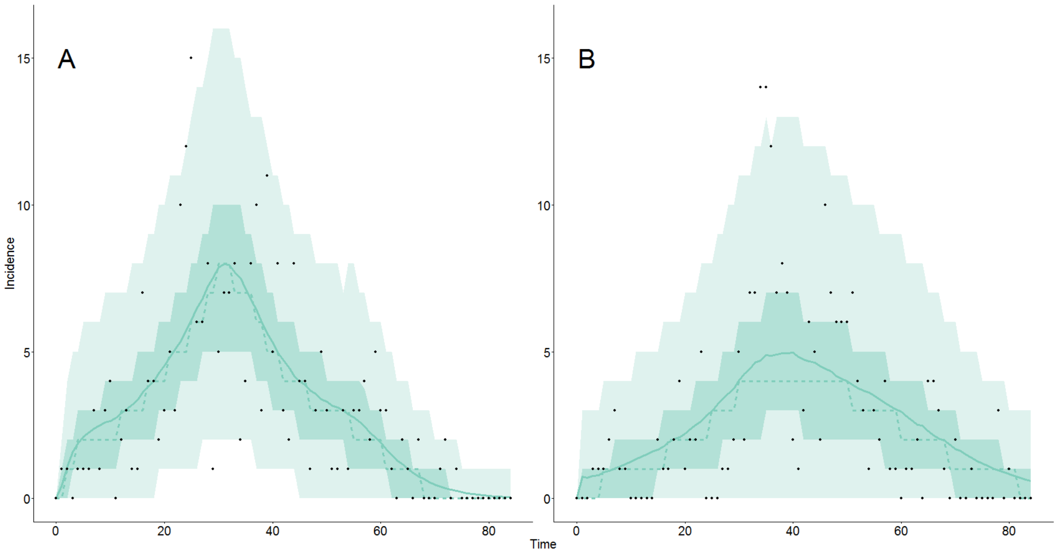

Figure S9: Comparison of our fitted model and observed daily incidence time series (black dots) reconstructed from the line list of EVD cases in Kikwit in 1995. A) Daily onset in the entire population ; B) Daily death in the entire population. The mean and median fits are represented by solid and dashed lines respectively. The dark and light shaded areas correspond to the 50% and 95% credible intervals.

#### S4.2 Parameter inference

MCMC chains were implemented using the *FitR* library in *R 3.2.3*. This library implements functions for model fitting and inference, providing the user with a Metropolis-Hastings algorithm, which adapts the covariance matrix during burn in to obtain an optimal acceptance rate and proposal distribution [9]. We simulated several chains to check they converge to the same stationary distribution. We ran these chains for 150,000 iterations, we then used 40000 iterations as burn in, with thinning (75%) to obtain the posterior distribution of parameters (Figure S10).

Table S3: Definition and estimation of the parameters of the deterministic model.

| Parameter | Description | Prior | Estimates : median (95% CI) |
|---|---|---|---|
| $\rho_{onset}$ | Reporting fraction of date of onset | Fixed (data) | 0.9 |
| $\rho_{death}$ | Reporting fraction of date of death | Fixed (data) | 0.95 |
| $\epsilon^{-1}$ | Incubation period | Fixed [10, 11] | 9.5 |
| $\gamma_{death}^{-1}$ | Duration onset to death | Fixed (data) | 10 |
| $\gamma_{recov}^{-1}$ | Duration onset to recovery | Fixed (data) | 18 |
| $\phi$ | Case-fatality Ratio | Fixed (data) | 0.78 |
| $N_c$ | Number of individuals in Community | Fixed [6] | 200,000 |
| $N_h$ | Number of HCW | Fixed [5, 7] | 900 |
| $T_0$ | Start of the study | Fixed (data) | April 07 |
| $\psi_{onset}$ | Overdispersion parameter for reporting of onset of symptoms | $\mathcal{U}[0-5]$ | 0.17 (0.03 - 0.41) |
| $\psi_{death}$ | Overdispersion parameter for reporting of dead cases | $\mathcal{U}[0-5]$ | 0.36 (0.13 - 0.73) |
| $\beta_{cc}$ | Community-to-community transmission parameter | $\mathcal{U}[0-100]$ | 0.06 (0.05-0.08) |
| $\beta_{ch}$ | Community-to-HCW transmission parameter | $\mathcal{U}[0-100]$ | 3.11 (1.00-7.10) |
| $\beta_{hn}$ | HCW-to-community transmission parameter | $\mathcal{U}[0-100]$ | 0.34 (0.21-0.54) |
| $\beta_{hh}$ | HCW-to-HCW transmission parameter | $\mathcal{U}[0-100]$ | 50.96 (34.86-76.24) |
| $\alpha_c$ | Shape of the change in community-to-community transmission | $\mathcal{U}[0-5]$ | 2.49 (0.21-4.86) |
| $\alpha_h$ | Shape of the change for HCW-related transmission | $\mathcal{U}[0-5]$ | 2.20 (0.23-4.83) |
| $\delta_c$ | Reduction of the within-community transmission rate following change of contact behaviour | $\mathcal{U}[0-5]$ | 2.90 (0.99-4.89) |
| $\delta_h$ | Reduction of the HCW-related transmission rates following change of contact behaviour | $\mathcal{U}[0-5]$ | 2.87 (1.09-4.84) |
| $T_h$ | Mid point date for the change of HCW behaviour | $\mathcal{U}[0-100]$ | 30 (27-35) days |
| $T_c$ | Mid point date for the change of community behaviour | $\mathcal{U}[0-115]$ | 55 (50-62) days |
| $E_{1cc}(T_0)$ | Number of exposed among community, infected by community member, at $T_0$ | $\mathcal{U}[0-115]$ | 13 (5 - 24) |
| $E_{1hc}(T_0)$ | Number of exposed among Community, infected by HCW, at $T_0$ | $\mathcal{U}[0-100]$ | 2 (0 - 8) |
| $E_{1ch}(T_0)$ | Number of exposed among HCW, infected by Community member, at $T_0$ | $\mathcal{U}[0-100]$ | 4 (1 - 10) |
| $E_{1hh}(T_0)$ | Number of exposed among HCW, infected by HCW, at $T_0$ | $\mathcal{U}[0-100]$ | 4 (0 - 12) |
| $I_c(T_0)$ | Number of infected among Community at $T_0$ | $\mathcal{U}[0-100]$ | 9 (1 - 24) |
| $I_h(T_0)$ | Number of infected among HCW at $T_0$ | $\mathcal{U}[0-100]$ | 2 (0 - 7) |
| $\tau_{pc}(t)$, $\tau_{bc}(t)$, $\tau_{ph}(t)$, $\tau_{bh}(t)$ | Vaccination rate for prime and boost vaccine per population | Fixed | 1,500 doses per day in total if vaccine campaign implemented |
| $\nu_p$, $\nu_b$ | Efficacy of prime and boost vaccine dose | Fixed | 0.8, 0.9 |

Shape parameters of the sigmoid functions have flat posterior distributions, from a certain value they cause an abrupt decrease of the transmission parameters. Also they are correlated with other parameters defining the sigmoid functions. We used non-informative priors for transmission rates, overdispersion, sigmoid function parameters, and the number of exposed and infected individuals at the start of the study period (Table S3).

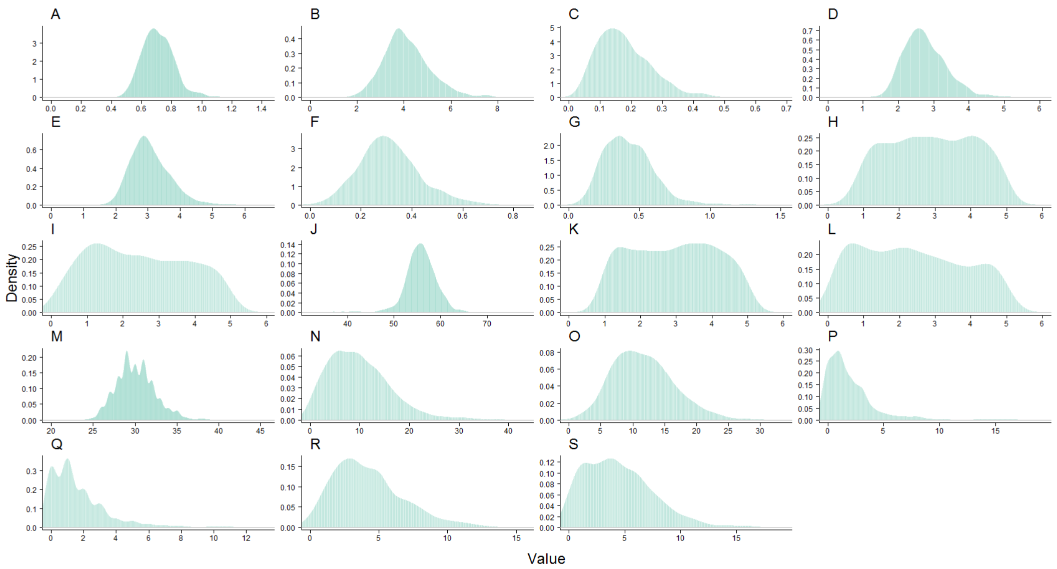

Figure S10: Posterior distribution of each one of the estimated parameters : A) $R_{0cc}$, B) $R_{0hc}$, C) $R_{0ch}$, D) $R_{0hh}$, E) $R_0$, F) $\psi_{onset}$, G) $\psi_{death}$, H) $\delta_c$, I )$\alpha_c$, J) $T_c$, K)$\delta_h$, L) $\alpha_h$, M) $T_h$, N) $I_{0c}$, O) $E_{0cc}$, P) $E_{0hc}$, Q) $I_{0h}$, R)$E_{0ch}$, S) $E_{0hh}$.

#### S4.3 Sensitivity to number of HCW in type 1 scenario

As there is uncertainty about the number of HCW in Kikwit in 1995, we tested the sensitivity of our model to this assumption. Thus we refitted the model with 700 or 1100 HCW, to compare with 900 in the main text. The log-likelihood values are the same for each chain (700 HCW : $-602.02$ ($-608.85$ - $-598.25$) 900 HCW : $-602.22$ ($-609.11$ - $-598.40$) 1100 HCW : $-602.34$ ($-609.17$ - $-598.83$)). The mean of the parameter distributions are similar for each chain (Figure S11). The estimation of the parameters of the model was therefore robust to changes in the number of HCW in Kikwit. We considered 900 HCW in the population as this value was supported by censuses.

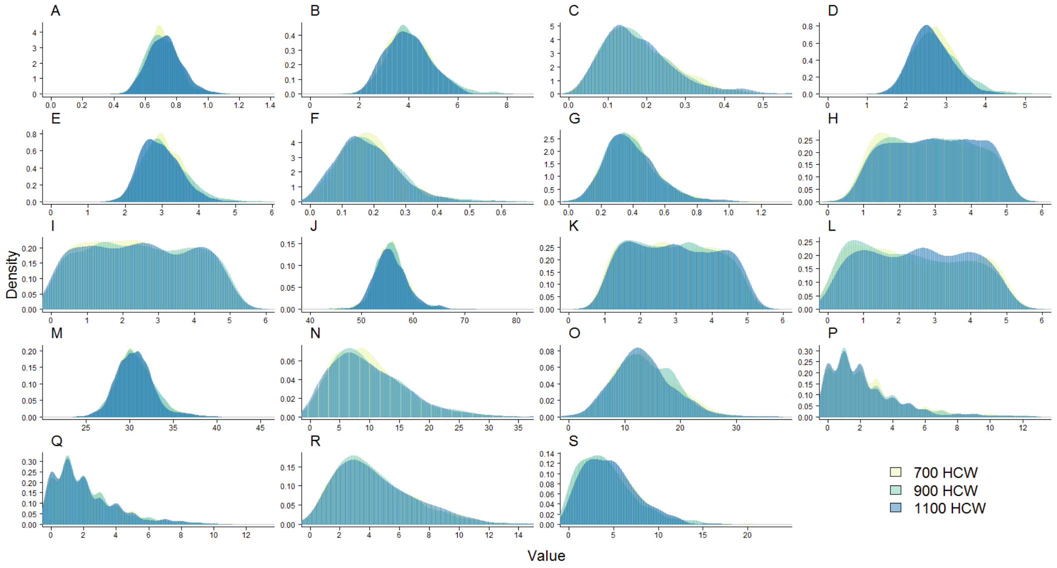

Figure S11: Comparison of the posterior distribution of each one of the estimated parameters depending on the number of HCW in Kikwit : A) $R_{0cc}$, B) $R_{0hc}$, C) $R_{0ch}$, D) $R_{0hh}$, E) $R_0$, F) $\psi_{onset}$, G) $\psi_{death}$, H) $\delta_c$, I) $\alpha_c$, J) $T_c$, K) $\delta_h$, L) $\alpha_h$, M) $T_h$, N) $I_{0c}$, O) $E_{0cc}$, P) $E_{0hc}$, Q) $I_{0h}$, R) $E_{0ch}$, S) $E_{0hh}$.

#### S4.4 Sensitivity to number of HCW in type 2 scenario

We used several features to describe type 2 outbreaks (Cumulative proportion of HCW among the total number of HCW infected through time, weekly proportion of HCW among the total number of HCW infected in EVD outbreaks through time, proportion of HCW infected through time, number of community cases ), we needed to show that the distributions didn't depend on the number of HCW in the population. Figure S12 shows that the number of HCW (700; 900 or 1,110) does not have a strong influence of the value of any of these parameters. Therefore we consider the results of our simulations able to describe settings with slightly different number of HCW in the population.

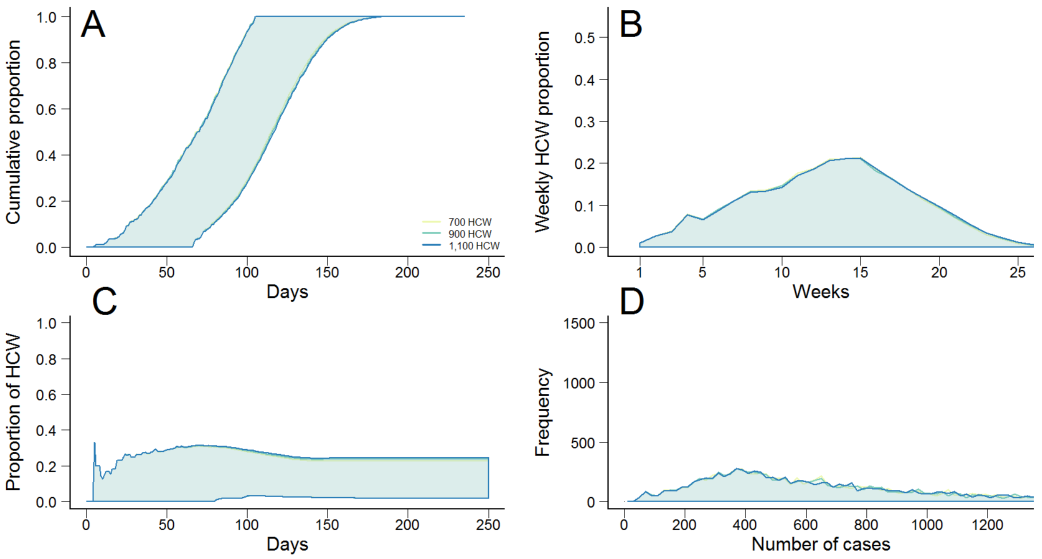

Figure S12: Comparison of the distribution of different features describing type 2 outbreaks, depending of the number of HCW included in the population. A) Cumulative proportion of HCW among the total number of HCW infected through time. B) Weekly proportion of HCW among the total number of HCW infected in EVD outbreaks through time. C) Proportion of HCW infected through time. D) Distribution of the number of community cases .

#### S4.5 Number of HCW initially infected in stochastic simulations

Vaccination reduced susceptibility to infection by $(1 - \nu_p)$ or $(1 - \nu_b)$. When we simulated an ahead-of-time vaccination of HCW, we modified the initial distribution of cases in type 1 outbreaks to illustrate the change in the susceptibility of individuals. Dependent on the value used for coverage, part of the HCW were already protected. We drew the number of exposed and infected HCW in this scenario using $E_{hh0}$, $E_{ch0}$ and $I_{h0}$ the number of exposed and infected HCW at $t = 0$ in the fitted parameter distributions, $\nu_p$ the efficacy of the vaccine and the value of coverage considered. The means were equal to $(1 - \nu_p)E_{0ch}(T_0)$, $(1 - \nu_p)E_{0hh}(T_0)$ and $(1 - \nu_p)I_h(T_0)$, multiplied by the value of coverage in each scenario. The distribution of initial distribution of HCW is represented in Figures S13 and S14. For vaccination scenarios including ahead-of-time vaccination of HCW in type 2 outbreaks, epidemics were seeded with 5 infected and 5 exposed community members, and no HCW infected.

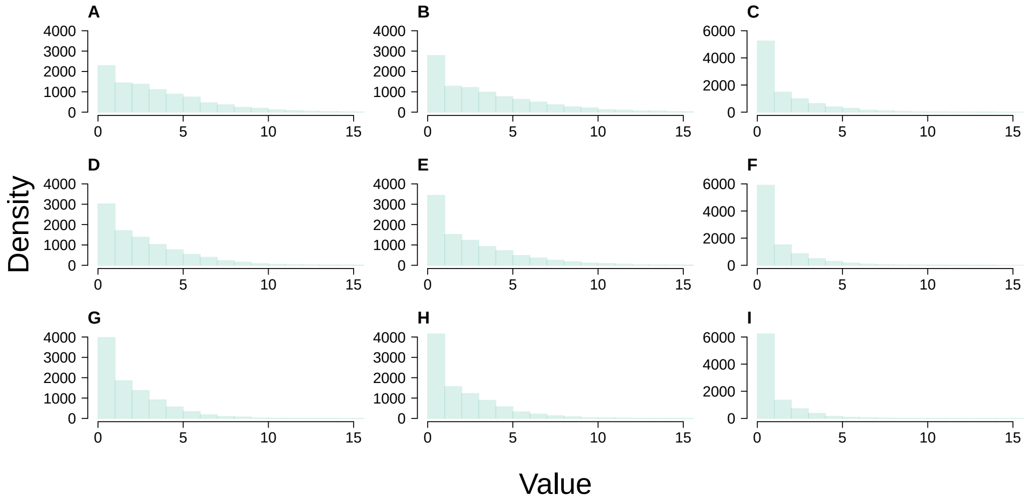

Figure S13: Number of HCW initially infected and exposed by Community members or other HCW in scenarios with ahead-of-time vaccination of HCW, stratified by coverage. A) Number of HCW initially exposed by Community members, B) by other HCW or C) infected with 10% coverage. D, E and F represent these values when coverage is 30% and G, H and I when half the HCW are vaccinated at $t = 0$

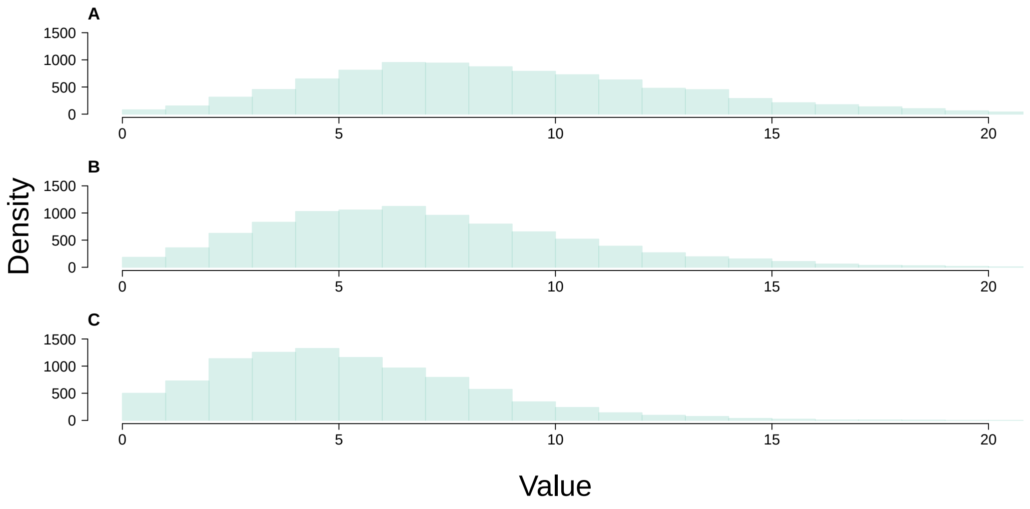

Figure S14: Total number of HCW exposed or infected when coverage is set to A) 10%, B) 30% and C)50%.

##### S4.6 Sensitivity to mid point date for the change of behaviour for type 2 scenario

We could not fit our model to the data and estimate parameters for the type 2 scenario. Therefore we analysed the impact of changing the mid point date for the change of behaviour on the dynamics of transmission among community members and HCW (Fig. S15). Decreasing or increasing the mid point date by 25 days did not change the proportion of HCW in the simulated outbreaks (Panels C and D), but greatly influenced the number of cases infected and the duration of outbreaks. The set of simulations where the mid point date is 125 days after the start of the outbreak was the closest to the data.

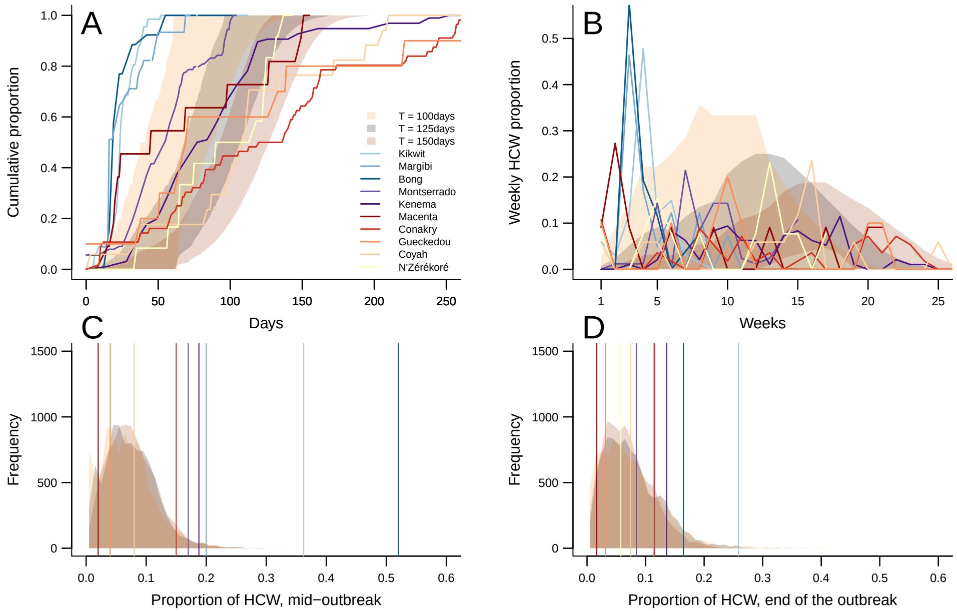

Figure S15: A) Cumulative proportion of HCW among the total number of HCW infected through time. B) Weekly proportion of HCW among the total number of HCW infected through time. C) Proportion of HCW among all cases infected at mid-outbreak. D) Proportion of HCW among all cases infected at the end of the outbreak Vertical lines in Figure C and D correspond to the value for each outbreak.

#### S5 Classification of epidemics

##### S5.1 Type 1 and 2 outbreaks

EVD outbreaks included in this study exhibited different transmission dynamics. Using a combination of factors, we classified these outbreaks into two types. We described each outbreak and disentangle the impact of HCW on the spread of the disease using various variables such as the total number of cases reported, the duration of outbreaks or the proportion of HCW infected through time (Figure S16). We did not include Kissidougou and Bomi in our analysis as less than 10 HCW were reported during these outbreaks. The distribution of the variables (Figure S17) showed clear differences between type 1 and type 2 outbreaks.

| | Total number of cases | Duration of outbreak | Duration first to last HCW infected | Time till 50 percent of infected HCW infected | Time till 80 percent of infected HCW infected | Percentage of HCW among infected at the end of the outbreak | Percentage of HCW among infected at halfway through by time | Difference in percent HCW - total vs halfway by time | Percentage of HCW among infected at halfway through by number of cases infected | Difference in percent HCW - total vs halfway by number of cases infected | Classification |
|---|---|---|---|---|---|---|---|---|---|---|---|
| Kikwit | 284 | 73 days | 45 days | 24 days | 33 days | 26% | 36% | 10% | 39% | 13% | 1 |
| Bong | 158 | 104 days | 39 days | 19 days | 32 days | 16% | 52% | 36% | 30% | 14% | 1 |
| Margibi | 394 | 72 days | 58 days | 16 days | 40 days | 11% | 20% | 9% | 17% | 6% | 1 |
| Montserrado | 1055 | 104 days | 104 days | 55 days | 81 days | 8% | 17% | 9% | 11% | 3% | 2 |
| Kenema | 706 | 250 days | 236 days | 77 days | 112 days | 14% | 18% | 4% | 19% | 5% | 2 |
| Conakry | 485 | 282 days | 260 days | 137 days | 174 days | 12% | 15% | 3% | 15% | 3% | 2 |
| Macenta | 660 | 155 days | 136 days | 45 days | 127 days | 2% | 2% | 0% | 2% | 0% | 2 |
| Gueckedou | 317 | 332 days | 303 days | 71 days | 221 days | 3% | 4% | 1% | 5% | 2% | 2 |
| Coyah | 228 | 324 days | 207 days | 109 days | 172 days | 7% | 8% | 1% | 6% | -1% | 2 |
| N'Zerekore | 229 | 168 days | 119 days | 118 days | 132 days | 6% | 4% | -2% | 4% | -2% | 2 |
| Kissidougou | 118 | 42 days | 39 days | 23 days | 41 days | 4% | 7% | 3% | 5% | 1% | too few HCW |
| Bomi | 80 | 90 days | 49 days | 16 days | 39 days | 8% | 8% | 0% | 11% | 3% | too few HCW |

Figure S16: Table describing the clustering of Ebola outbreaks using different features.

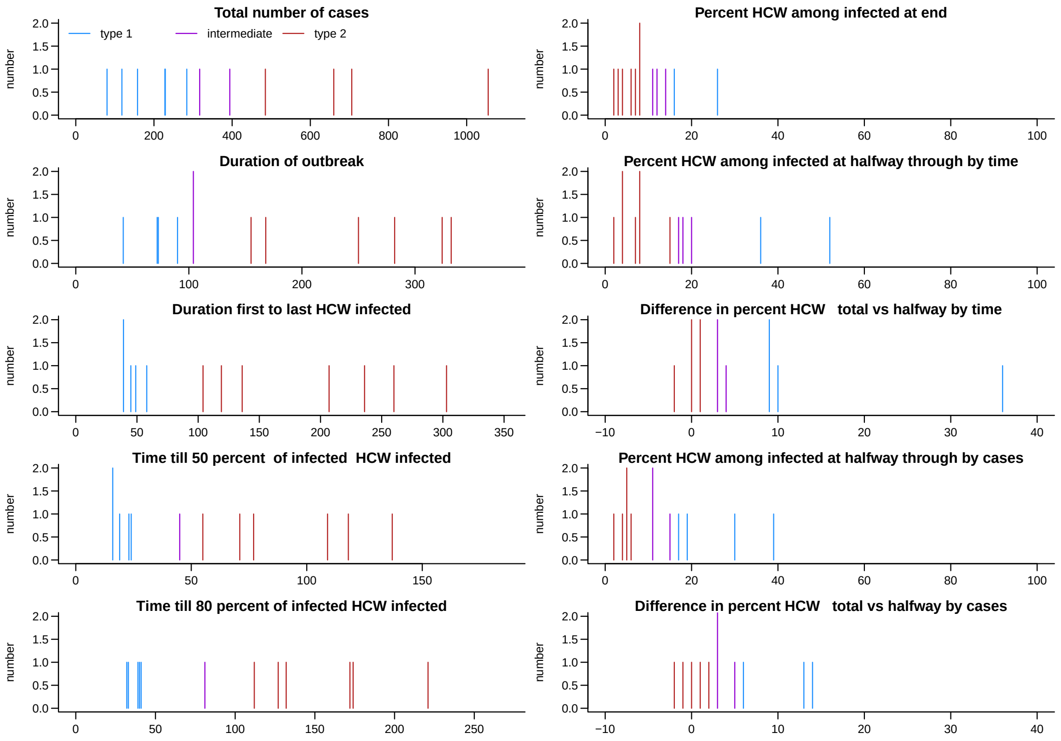

Figure S17: Distribution of the values described in Figure S16 used to cluster EVD outbreaks

In Kikwit, DRC, Kissidougou, Guinea and Bong, Bomi and Margibi counties, Liberia, HCW tended to be infected early in the outbreak, and few subsequent community transmissions were observed later on. We found evidence that in Kikwit, a type 1 outbreak, HCW caused high number of cases among Community and HCW. In contrast, in Kenema, Montserrado, Macenta,N'Zerekore, Gueckedou and Conakry the increase in HCW infections was slower, with a more stable number of weekly infections among HCW until the end of the outbreak (Figure S18). In this latter setting, the number of infected cases grew bigger without important HCW transmission, showing sustainable transmission among community members (i.e. reproduction number is greater than 1).

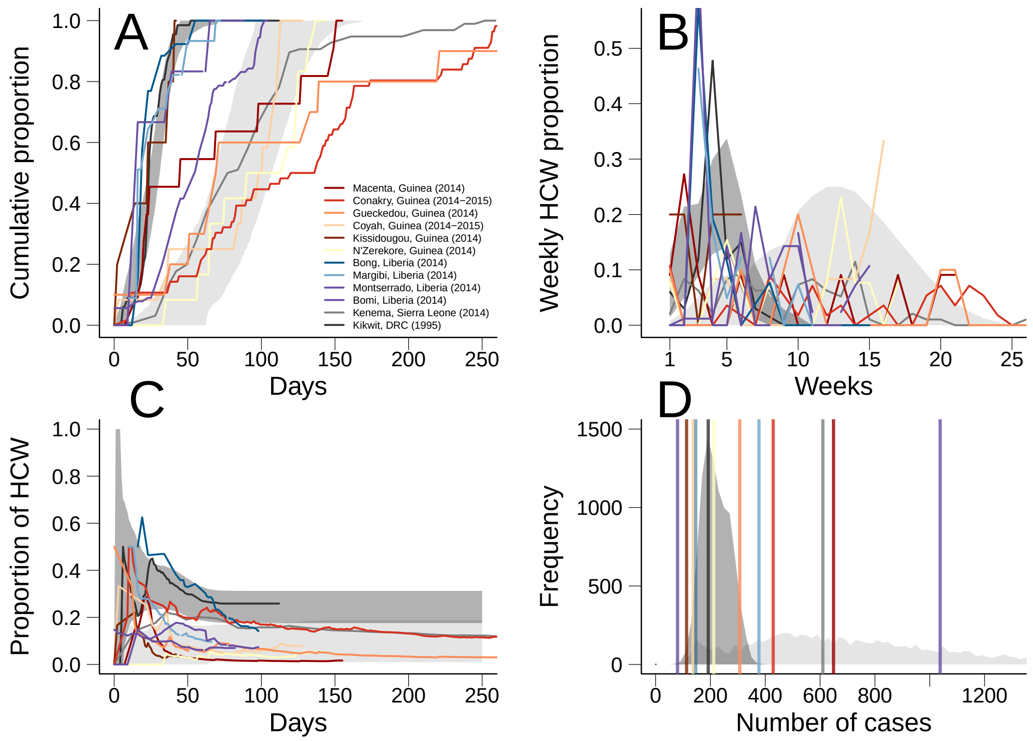

Figure S18: Classification of EVD outbreaks into two types according to their HCW-related dynamics. Dark grey marks the 95% CI in type 1 outbreak simulations, light grey areas correspond to 95% CI for type 2 outbreak simulations. A) Cumulative proportion of HCW among the total number of HCW infected through time. B) Weekly proportion of HCW among the total number of HCW infected in EVD outbreaks through time. C) Proportion of HCW infected through time. D) Distribution of the number of community cases, where vertical lines represent the number of community cases observed in each outbreak.

In type 2 scenario, as fewer people were infected initially and transmission lasted longer, the range of number of cases per outbreak was broader than the type 1 scenario. The proportion of HCW among infected in the simulations matched the proportions observed in the outbreaks from the second cluster (Fig. S19). The type 2 scenario generated outbreaks with larger number of cases (Figure S19), which corresponded to the outbreaks in the second clusters. The cumulative proportion of HCW infected over time in the simulations highlights a much more linear increase, which was observed in West Africa (e.g. in Kenema) (Fig S19). The Figure S20 highlights the differences between the two clusters of Ebola outbreaks: high percentage of HCW infected are reached in much shorter time in Kikwit and Bong district than in the other outbreaks, this characteristic is well reproduced by the two simulation scenarios.

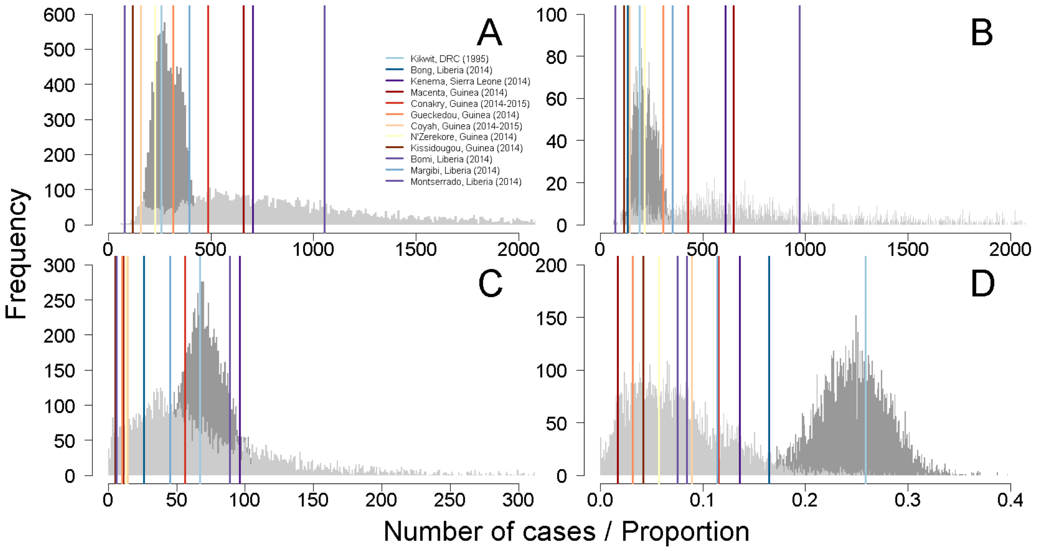

Figure S19: Distribution of cases from simulations. Dark grey is type 1 outbreak simulation, light grey is type 2 outbreak simulation. The coloured line marks the observed value from the example outbreaks. A) Total number of cases per outbreak, B) Number of cases in the community, C) Number of HCW infected, D) Proportion of Heath Care Workers.

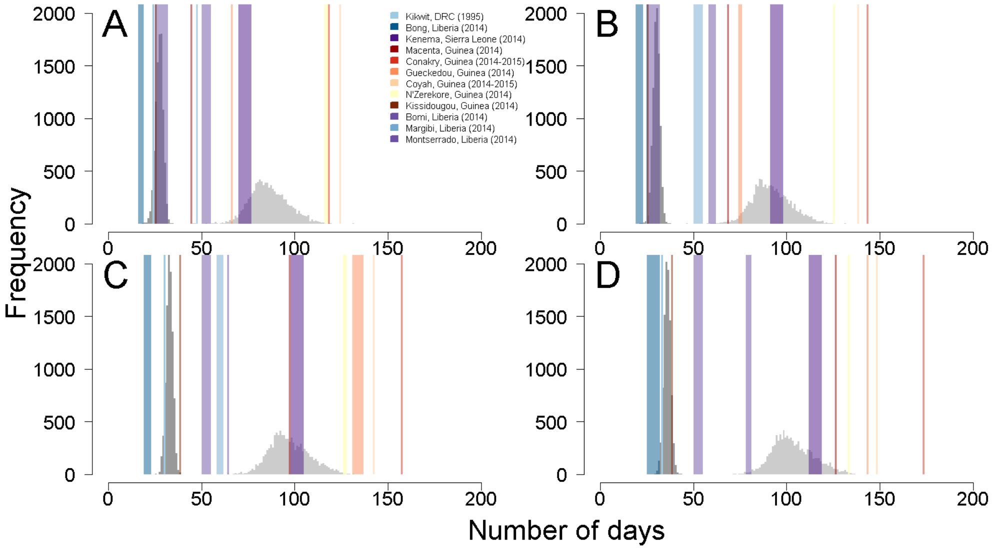

Figure S20: Number of days after the start of the outbreak before reaching A) 50% ; B) 60% ; C) 70% ; D) 80% of the total number of HCW infected. Dark grey is type 1 outbreak simulation, light grey is type 2 outbreak simulation. The coloured window marks the value in the example outbreaks, where a window indicates uncertainty in the true value for that outbreak, ie. 1 day, or 1 week window.

#### S5.2 Comparison of reproduction numbers in type 1 and type 2 scenarios

The Figure S21 shows the values of the overall Reproduction Number through time in each scenario. We generated the type 2 scenario to reproduce outbreaks with linear increase of cases over a long period of time, therefore the drop in the reproduction number is smoother, and transmission is sustainable during several months.

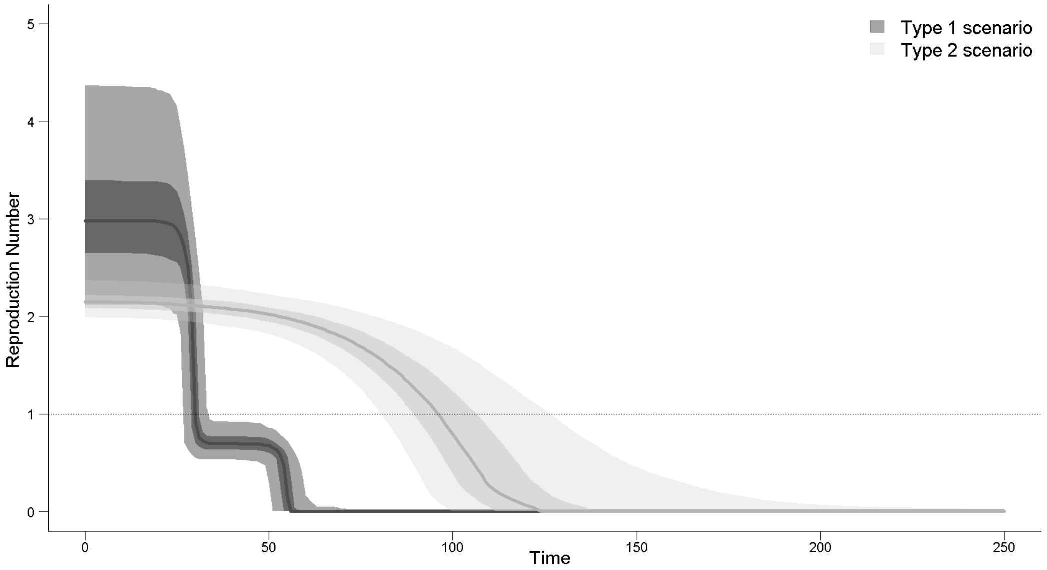

Figure S21: Comparison of the mean reproduction number trajectories used for the two scenarios, with 50% (dark) and 95% CI (light). Type 1 outbreaks (red) use parameters fitted to data from Kikwit, DRC, and type 2 outbreaks (blue) use a modelled scenario described in Methods.

### S6 Effect of Vaccination

#### S6.1 Proportion of cases averted

The proportion of remaining cases compared with the baseline simulation without vaccination is displayed in Figure S22. In the type 1 scenario, the indirect effect of ahead-of-time vaccination of HCW when high (50%) coverage was reached (vaccination strategy e), decreased the total number of cases. The combined vaccination strategies (f, g, h) were very comparable to the ahead-of-time HCW in the type 1 scenario. In contrast, in the type 2 scenario, there was a large difference between ahead-of-time HCW vaccination only (strategies c,d

and e), and the corresponding combined approaches. This occurred because of the sustained within-community transmission in the type 2 scenario.
Reactive mass vaccination strategies (green boxplots) substantially reduced the number of cases in the population in the type 2 scenario. In the type 1 scenario, the overall reproduction number got below 1 faster, as control measures and behaviour changes occurred early and struggled down the spread of EVD, thus reactive mass vaccination had very little impact.

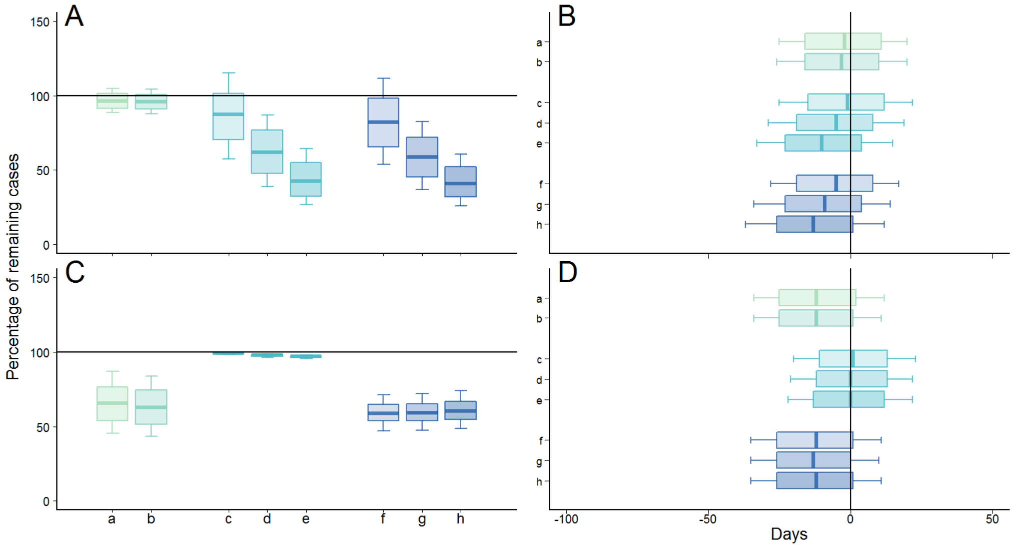

Figure S22: Simulation results comparing to the baseline simulation without vaccination. A) Percentage of remaining cases duration of the outbreak in the type 1 scenario; B) Comparison of the duration of the outbreak in the type 1 scenario; C) Percentage of remaining cases in the type 2 scenario ; D) Comparison of the duration of the outbreak in the type 2 scenario. Each subscript represents a vaccination strategy: reactive mass vaccination only, with (a) prime-boost vaccine or (b) single dose vaccine; ahead-of-time HCW vaccination only, with (c) 10% coverage, (d) 30% coverage or (e) 50% coverage among HCW; reactive mass vaccination combined with ahead-of-time HCW vaccination, with (f) 10% coverage, (g) 30% coverage or (h) 50% coverage among HCW. Rectangles represent the 50% CI, the arrows show the 75% CI

#### S6.2 Effect of vaccination on each occupation group

The distributions of the number of cases in each occupation group show that the vaccination regimes had different impact in each occupation group (Figure S23). In the type 1 scenario, where HCW "catalyse" the epidemic, reactive mass vaccination campaigns alone were unable to reduce the number of HCW infected. This was caused by the abrupt and early decrease in HCW-related transmission parameters. Furthermore, as transmission among community members was not sustainable, the spread among community members dropped quickly after, leading to a poor added effect of mass vaccination campaigns. On the other hand, ahead-of-time vaccination of HCW was able to reduce the number of community members infected in both scenarios, due to indirect effects, which depended on the coverage of HCW. In the type 2 scenario, reactive mass vaccination campaigns diminished the number of HCW infected, because HCW were prioritised for vaccination in the model. The prolonged sustainable transmission in the community was cut off by the mass vaccination campaign, leading to an important drop in the number of community members infected.

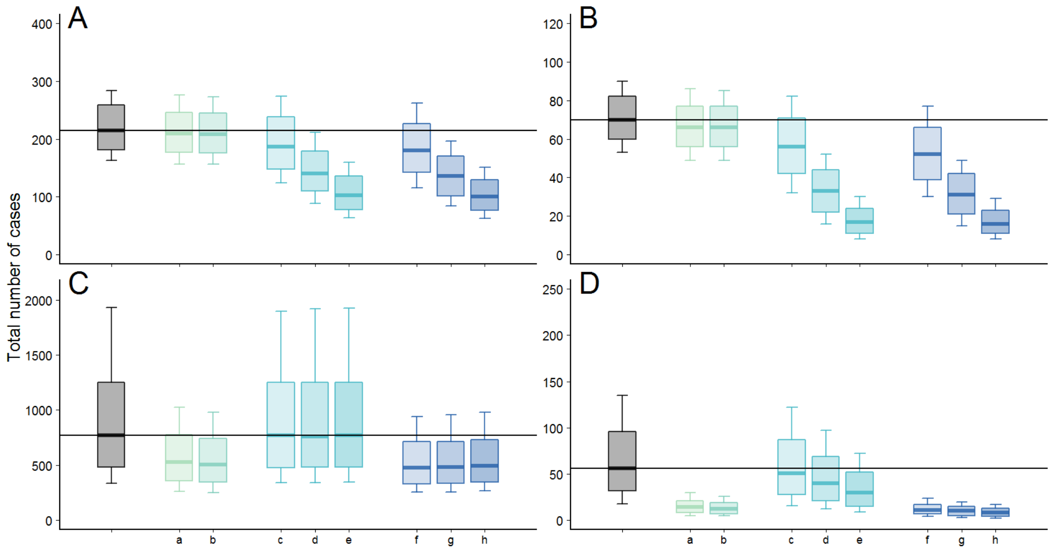

Figure S23: Number of cases in the stochastic simulations for each scenario, stratified by occupation group. A) Number of cases among community, type 1 scenario ; B) Number of cases among HCW, type 1 scenario; C) Number of cases among community, type 2 scenario ; D) Number of cases among HCW, type 2 scenario. Each subscript represents a vaccination strategy: reactive mass vaccination only, with (a) prime-boost vaccine or (b) single dose vaccine; ahead-of-time HCW vaccination only, with (c) 10% coverage, (d) 30% coverage or (e) 50% coverage among HCW; reactive mass vaccination combined with ahead-of-time HCW vaccination, with (f) 10% coverage, (g) 30% coverage or (h) 50% coverage among HCW.

##### S6.3 Comparison of 10%, 30% and 50% coverage of HCW

The indirect effect caused by ahead-of-time HCW vaccination was noticeable with 10% coverage (Figure S24) in the type 1 scenario. We observed that with 50% coverage (Figure S26), the daily incidence was substantially reduced for both scenarios, but the outbreak persisted for a longer time, which could potentially increase the risk of spread to other areas.

Reactive mass vaccination caused a drop in the the daily incidence. In the type 1 scenario the reactive mass vaccination strategies sped up the decrease in number of cases after the peak. In the type 2 scenario, the reactive mass vaccination strategy resulted in fewer cases than the ahead-of-time HCW vaccination only, even with 50% coverage.

The figures S24, S25 and S26 show that in case of large outbreaks happening when ahead-of-time HCW vaccination is already implemented, starting a reactive mass vaccination campaign (combined approach) leads to a plateau in the daily incidence, and speeds up the reduction of transmission.

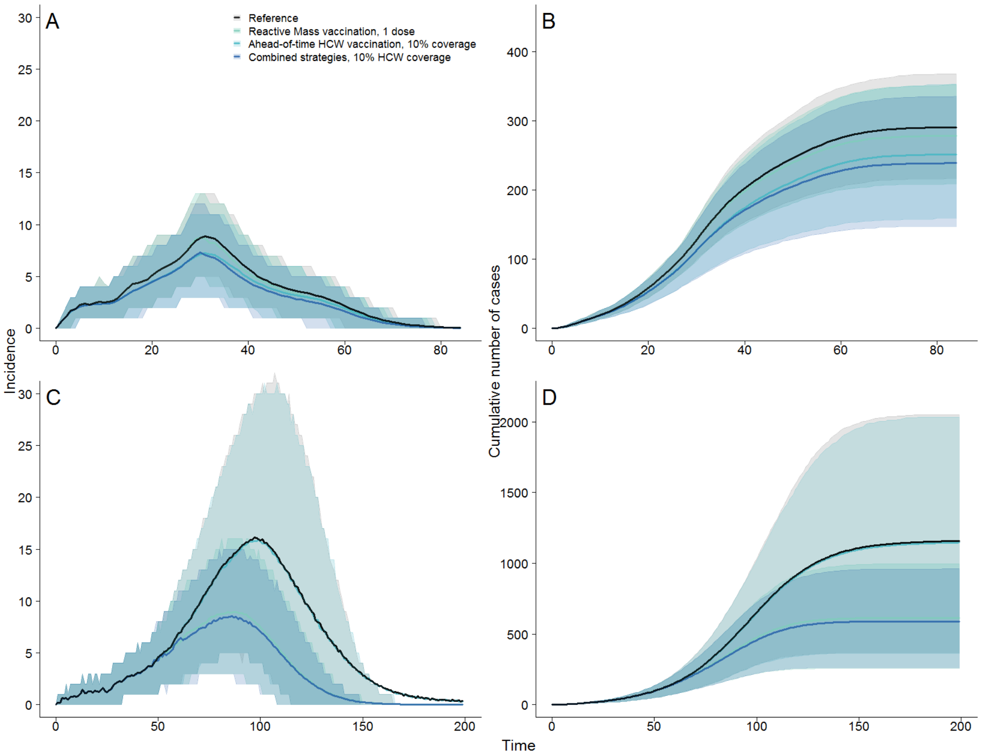

Figure S24: Simulated epidemics under three vaccination scenarios. A and B show incidence and cumulative number of cases in the type 1 scenario, for HCW and community members combined. C and D show under the type 2 scenario. The vaccination strategies compared are : Reactive mass vaccination campaign with single-dose vaccine, ahead-of-time vaccination campaign among HCW with 10% coverage and combined strategies with 10% coverage among HCW. Note different y axis. The shaded areas show the 75%CIs. Plain lines represent the mean of the daily number of cases in the simulation set.

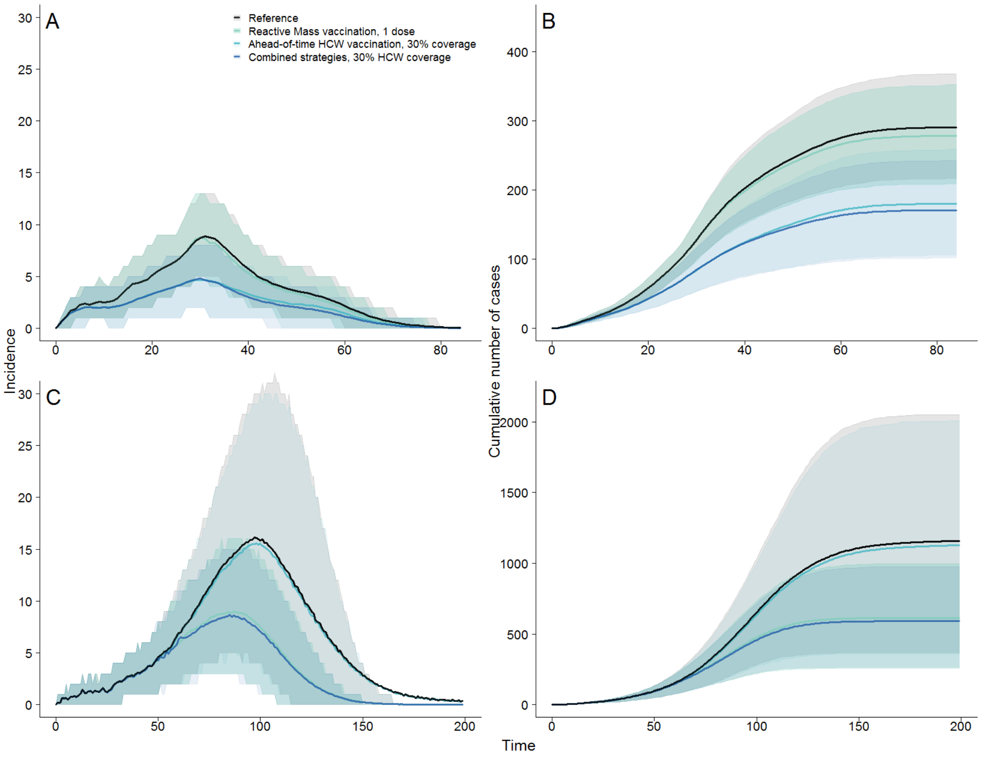

Figure S25: Simulated epidemics under three vaccination scenarios. A and B show incidence and cumulative number of cases in the type 1 scenario, for HCW and community members combined. C and D show under the type 2 scenario. The vaccination strategies compared are : Reactive mass vaccination campaign with single-dose vaccine, ahead-of-time vaccination campaign among HCW with 30% coverage and combined strategies with 30% coverage among HCW. Note different y axis. The shaded areas show the 75%CIs. Plain lines represent the mean of the daily number of cases in the simulation set.

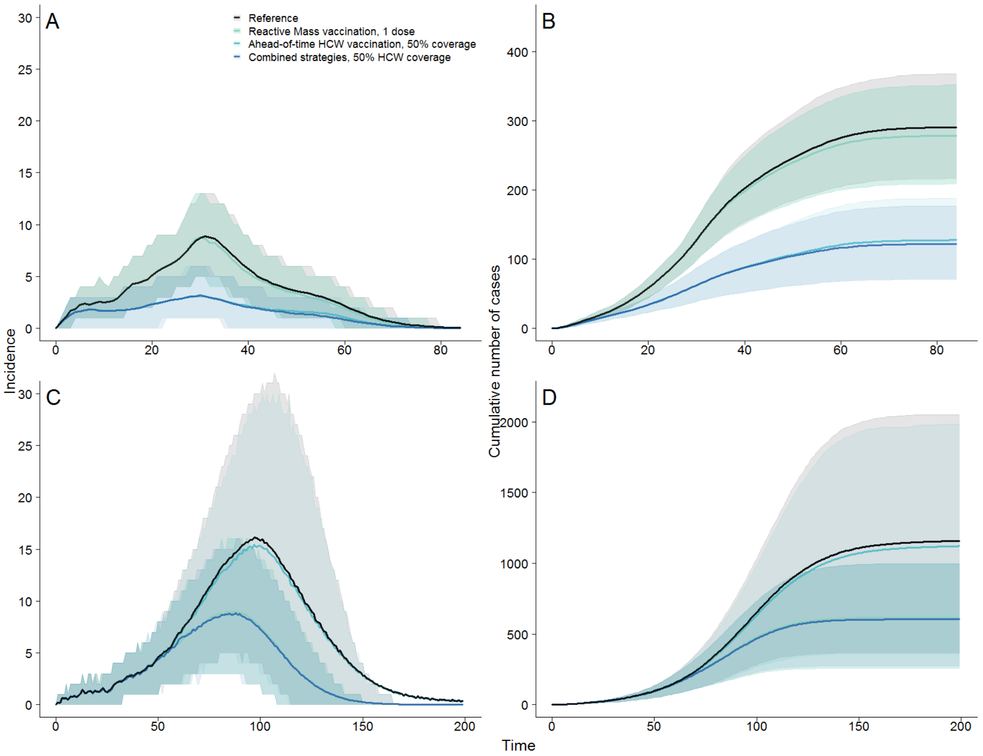

Figure S26: Simulated epidemics under three vaccination scenarios. A and B show incidence and cumulative number of cases in the type 1 scenario, for HCW and community members combined. C and D show under the type 2 scenario. The vaccination strategies compared are : Reactive mass vaccination campaign with single-dose vaccine, ahead-of-time vaccination campaign among HCW with 50% coverage and combined strategies with 50% coverage among HCW. Note different y axis. The shaded areas show the 75%CIs. Plain lines represent the mean of the daily number of cases in the simulation set.

#### S6.4 Simulated trajectories for comparison between coverage levels

Figure S27 shows the impact of different levels of coverage on daily incidence, and cumulative number of cases, as well as a comparison of single-dose and prime-boost vaccination, for the type 1 and type 2 scenarios.
The prime-boost vaccine induces a later drop than the single-dose vaccine due to the two week delay between the doses.
The variation in coverage levels causes marginal differences in epidemic trajectory, and graded decrease in number of cases as the coverage increases. A larger difference between coverage levels is seen in the type 1 scenario, where HCW "catalyse" the outbreak early on, and therefore their vaccine coverage level is critical to the size of the outbreak.

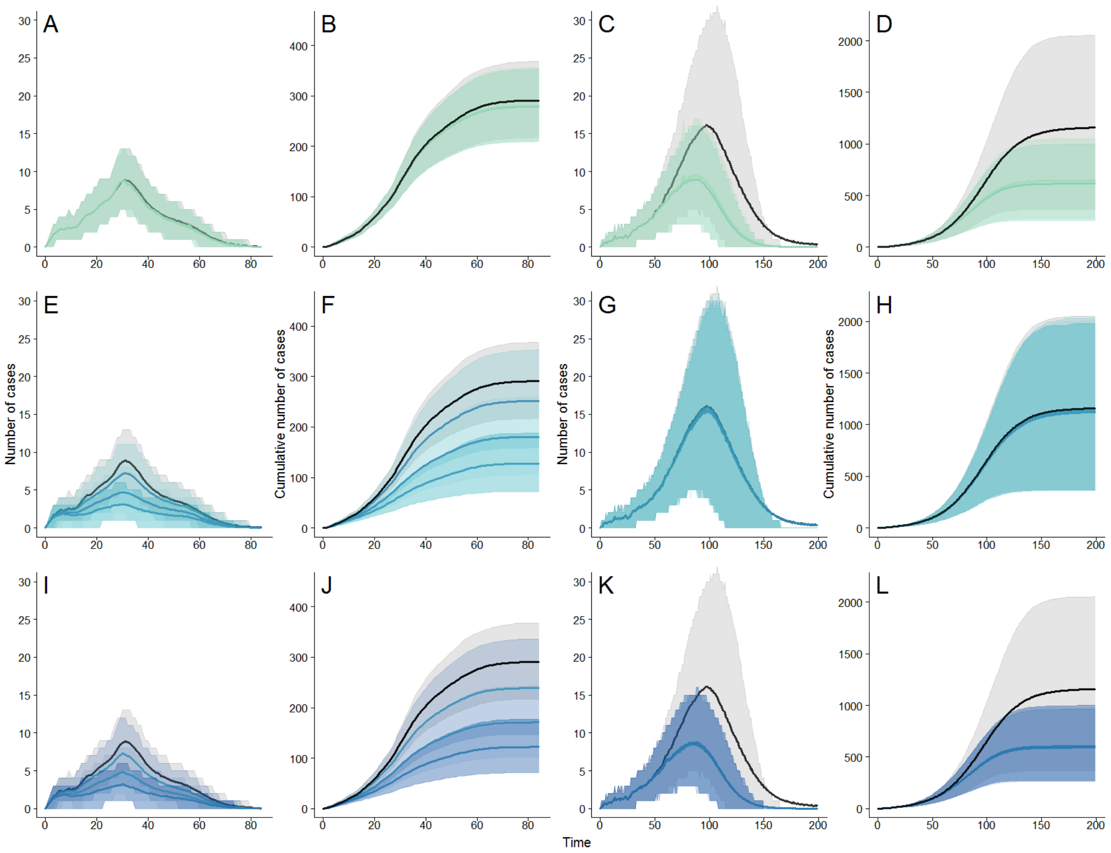

Figure S27: Trajectories of simulated epidemics. The shaded areas show the 75% credibility intervals. The plain lines represent the mean of the daily number of cases in the set. A-B, E-F and I-J show incidence and cumulative number of cases in the type 1 scenario, for HCW and community members combined. C-D, G-H and K-L show under the type 2 scenario. The reactive mass vaccination strategies with one (dark green) and two (light green) doses are compared to the baseline in A to D. The ahead-of-time HCW vaccination campaigns only with 10%, 30% and 50% coverage are compared to the reference in E to H. The combined campaigns with 10%, 30% and 50% coverage are compared to the reference in I to L.

#### S6.5 Number of cases per scenario

The tables S4-S5 show the number of cases generated and the durations of outbreaks in the stochastic simulations.

| Strategy | Total number of cases | Community cases | HCW infected | Duration |
|---|---|---|---|---|
| baseline | 287 [180-408] | 215 [133-321] | 70 [40-106] | 115 [92-154] |
| a | 278 [169-403] | 209 [125-318] | 66 [37-102] | 113 [92-150] |
| b | 277 [168-401] | 208 [124-315] | 66 [37-101] | 112 [92-150] |
| c | 245 [100-430] | 187 [81-335] | 56 [18-102] | 114 [90-153] |
| d | 174 [69-326] | 140 [59-266] | 33 [8-65] | 110 [87-151] |
| e | 121 [50-243] | 103 [45-208] | 17 [3-40] | 106 [82-147] |
| f | 233 [96-414] | 180 [76-327] | 52 [16-98] | 110 [88-147] |
| g | 167 [64-307] | 136 [56-252] | 31 [7-62] | 106 [82-144] |
| h | 117 [48-223] | 100 [43-190] | 16 [3-38] | 103 [78-141] |

Table S4: Number of cases depending on the vaccination strategy implemented in the simulations in scenario 1. The median is given for each setting, with the 95% CI. Duration is in days.

| Strategy | Total number of cases | Community cases | HCW infected | Duration |
|---|---|---|---|---|
| baseline | 839 [170-4305] | 767 [156-4158] | 56 [5-263] | 184 [149-229] |
| a | 546 [159-1764] | 528 [153-1722] | 14 [2-52] | 174 [141-223] |
| b | 521 [157-1674] | 506 [152-1650] | 12 [1-43] | 173 [140-223] |
| c | 834 [163-4396] | 769 [154-4219] | 51 [5-232] | 186 [149-250] |
| d | 812 [163-4225] | 760 [157-4139] | 40 [3-184] | 185 [148-251] |
| e | 813 [163-4245] | 772 [156-4200] | 30 [3-130] | 185 [148-252] |
| f | 491 [145-1635] | 478 [142-1607] | 11 [1-40] | 173 [139-223] |
| g | 496 [148-1643] | 484 [144-1620] | 10 [1-34] | 173 [139-223] |
| h | 504 [162-1663] | 495 [157-1638] | 8 [1-29] | 173 [139-224] |

Table S5: Number of cases depending on the vaccination strategy implemented in the simulations in scenario 2. The median is given for each setting, with the 95& CI. Duration is in days.

### S7 Sensitivity analysis on HCW transmission in type 2 scenario

The data from type 2 outbreaks were too incomplete to fit the model, we decided to use parameter sets from previous studies. We also explored other parameter distributions that could generate simulation with similar dynamics y generating an alternative type 2 scenario as a sensitivity analysis (Fig. S28, S29). In this scenario, HCW have similar transmission characteristics as community members.

| | | HCW to HCW | Community to HCW | HCW to community | Community to community | Overall |
|---|---|---|---|---|---|---|
| Type 1 scenario | $R_0$ | 2.68 (1.83-4.01) | 0.16 (0.05-0.37) | 3.99 (2.47-6.29) | 0.70 (0.53-0.95) | 2.98 (2.11-4.36) |
| | $T_{change}$ *(days)* | $T_h$ = 30 (27-35) | $T_h$ | $T_h$ | $T_c$ = 55 (50-62) | |
| | shape | $\alpha_h$ = 2.20 (0.23-4.83) | $\alpha_h$ | $\alpha_h$ | $\alpha_c$ = 2.49 (0.21-4.86) | |
| Type 2 scenario | $R_0$ | 1.50 (1.33-1.75) | 0.16 (0.05-0.37) | 1.50 (1.33-1.75) | 1.50 (1.33-1.75) | 1.99 (1.71-2.38) |
| | $T_{change}$ *(days)* | $T_h$ = 130 | $T_h$ | $T_h$ | $T_c$ = 155 (148-163) | |
| | shape | $\alpha_c$ = 0.05 | $\alpha_c$ | $\alpha_c$ | $\alpha_c$ | |

Figure S28: Parameters to generate alternative type 2 simulations.

The features of the outbreaks generated with the sensitivity analysis scenario simulations are shown in Figure S30, along with the data. We observed the dynamics of the generated outbreaks was similar to the main scenario 2 and to the data. The bottom panels of Figure S31 shows the impact of each control measures on the number of cases in the community and among HCW in alternative type 2 simulations. The strategies had similar effects on main and alternative type 2 outbreaks and led to similar conclusions: reactive mass vaccination strategies had an important impact on both HCW and community members, and ahead of time vaccination of HCW had limited impact on community members in both type 2 scenarios.

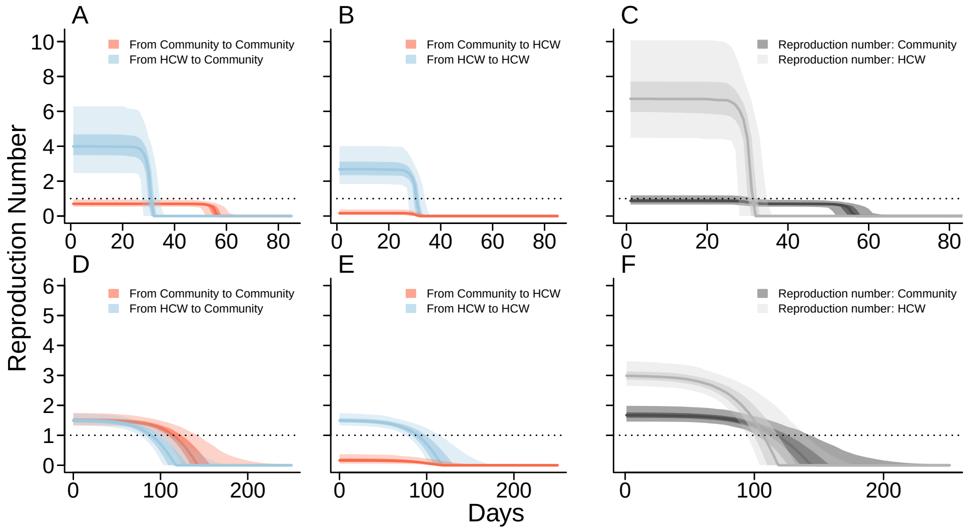

Figure S29: A and D) HCW-to-community reproduction number (blue) and community-to-community (red) are given with mean and 50% and 95% CI. B and E) HCW-to-HCW reproduction number (blue) and community-to-HCW reproduction number (red) decrease at the same time, Th. The horizontal line indicates R=1. C and F) The overall reproduction number trajectories for Community members (dark grey) and HCW (light grey), with 50% and 95% CI.

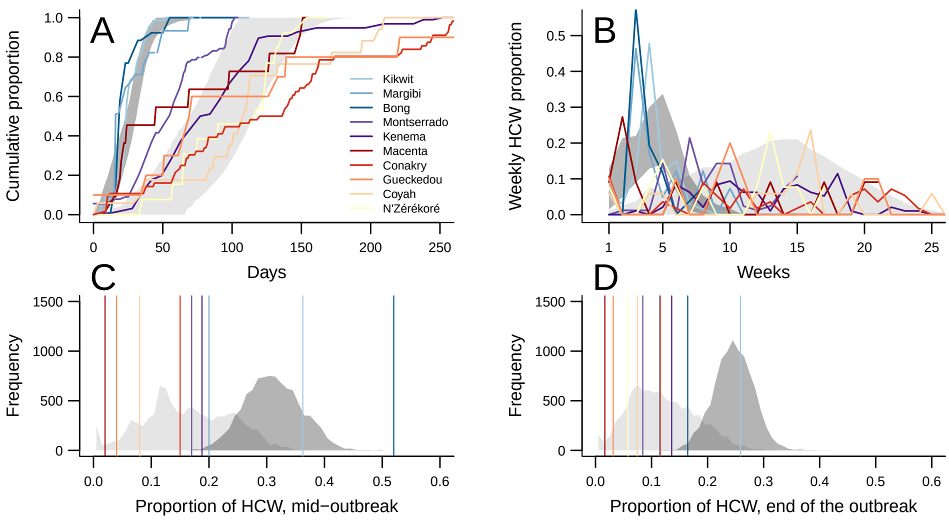

Figure S30: Dark grey marks the 95% CI in type 1 outbreak simulations, light grey areas correspond to 95% CI for type 2 outbreak simulations. Each colour corresponds to an outbreak, we used blue shades for type 1 outbreaks and red/purple shades for type 2. A) Cumulative proportion of HCW among the total number of HCW infected through time. B) Weekly proportion of HCW among the total number of HCW infected through time. C) Proportion of HCW among all cases infected at mid-outbreak. D) Proportion of HCW among all cases infected at the end of the outbreak Vertical lines in Figure C and D correspond to the value for each outbreak.

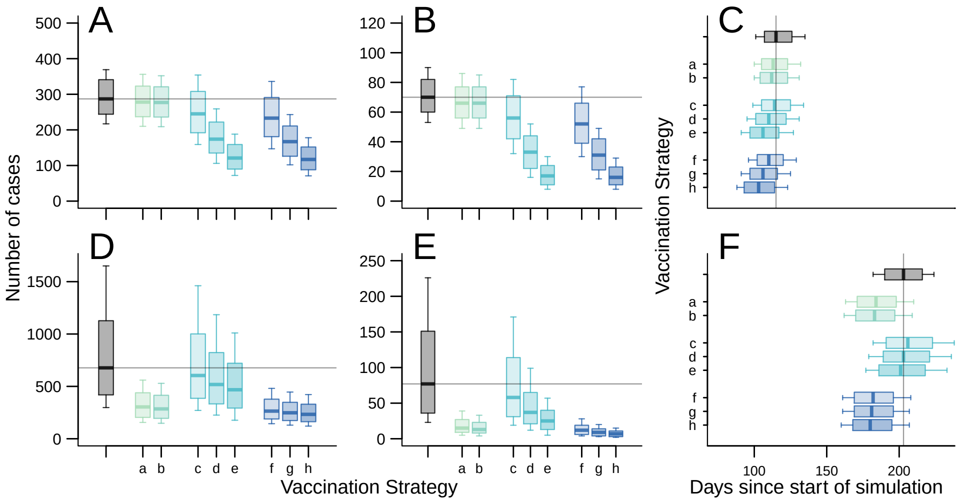

Figure S31: In type 1 outbreaks A) Number of cases in the entire population, B) number of cases in the 900 simulated HCW, and C) time to extinction; In type 2 outbreaks the: D) Number of cases in the entire population, E) number of cases in the 900 simulated HCW, and F) time to extinction. Boxplots show median value, rectangles mark 50% CI, and whiskers the 75% CI. Simulations without vaccination are shown in grey, and each colour represents a vaccination strategy (Table 3): reactive mass vaccination with (a) prime-boost vaccine or (b) single dose vaccine; ahead-of-time HCW vaccination only, with coverage in HCW of (c) 10%, (d) 30% or (e) 50%; ahead-of-time HCW vaccination plus reactive mass vaccination, with coverage in HCW of (f) 10%, (g) 30% or (h) 50%. We give the 75% CI due to high variation in the simulation sets, and 95% CI are given in the Supplement (S6.5) Note different y-axes.

### References

[1] World Health Organization. Health worker Ebola infections in Guinea, Liberia and Sierra Leone - A preliminary report. (May):1–16, 2015.

[2] Caitlin Rivers. Data for the 2014 Ebola Outbreak in West Africa, 2014.

[3] Mikiko Senga, Kimberly Pringle, Andrew Ramsay, David M. Brett-Major, Robert A. Fowler, Issa French, Mohamed Vandi, Josephine Sellu, Christian Pratt, Josephine Saidu, Nahoko Shindo, and Daniel G. Bausch. Factors underlying Ebola virus infection among health workers, Kenema, Sierra Leone, 2014-2015. *Clinical Infectious Diseases*, 63(4):454–459, 2016.

[4] J J Muyembe-Tamfum, M Kipasa, C Kiyungu, and R Colebunders. Ebola outbreak in Kikwit, Democratic Republic of the Congo: discovery and control measures. *The Journal of infectious diseases*, 179 Suppl(Suppl 1):S259–S262, 1999.

[5] Oyewale Tomori, Jeanne Bertolli, Pierre E Rollin, Yon Fleerackers, Yves Guimard, Ann De Roo, Heinz Feldmann, Felicity Burt, Robert Swanepoel, Scott Killian, Ali S Khan, Kweteminga Tshioko, Mpia Bwaka, Roger Ndambe, C J Peters, and Thomas G Ksiazek. Serologic Survey among Hospital and Health Center Workers during the Ebola Hemorrhagic Fever Outbreak in Kikwit, Democratic Republic of the Congo, 1995 formed by coating polyvinyl chloride microtiter plates overnight. *The Journal of Infectious Diseases*, 179(Suppl 1):98–101, 1999.

[6] S F Dowell, R Mukunu, T G Ksiazek, a S Khan, P E Rollin, and C J Peters. Transmission of Ebola hemorrhagic fever: a study of risk factors in family members, Kikwit, Democratic Republic of the Congo, 1995. Commission de Lutte contre les Epidémies à Kikwit. *The Journal of infectious diseases*, 179 Suppl:S87–S91, 1999.

[7] Unité de pilotage du DSRP Ministère du Plan RDC. Monographie De La Province Du Nord- Kivu. (Draft 4):1–134, 2005.

[8] O. Diekmann, J. A P Heesterbeek, and J. A J Metz. On the definition and the computation of the basic reproduction ratio R0 in models for infectious diseases in heterogeneous populations. *Journal of Mathematical Biology*, 28(4):365–382, 1990.

[9] Gareth O Roberts and Jeffrey S Rosenthal. Examples of adaptive MCMC. *Journal of Computational and Graphical Statistics*, 18(2):349–367, 2009.

[10] World Health Organization. Ebola virus disease. *World Health Organization*, page 1, 2015.

[11] J Legrand, R F Grais, P Y Boelle, A J Valleron, and A Flahault. Understanding the dynamics of Ebola epidemics. *Epidemiology and infection*, 135(4):610–21, 2007.
